## Supplementary Figures for "White matter myelination during early infancy is explained by spatial gradients and myelin content at birth"

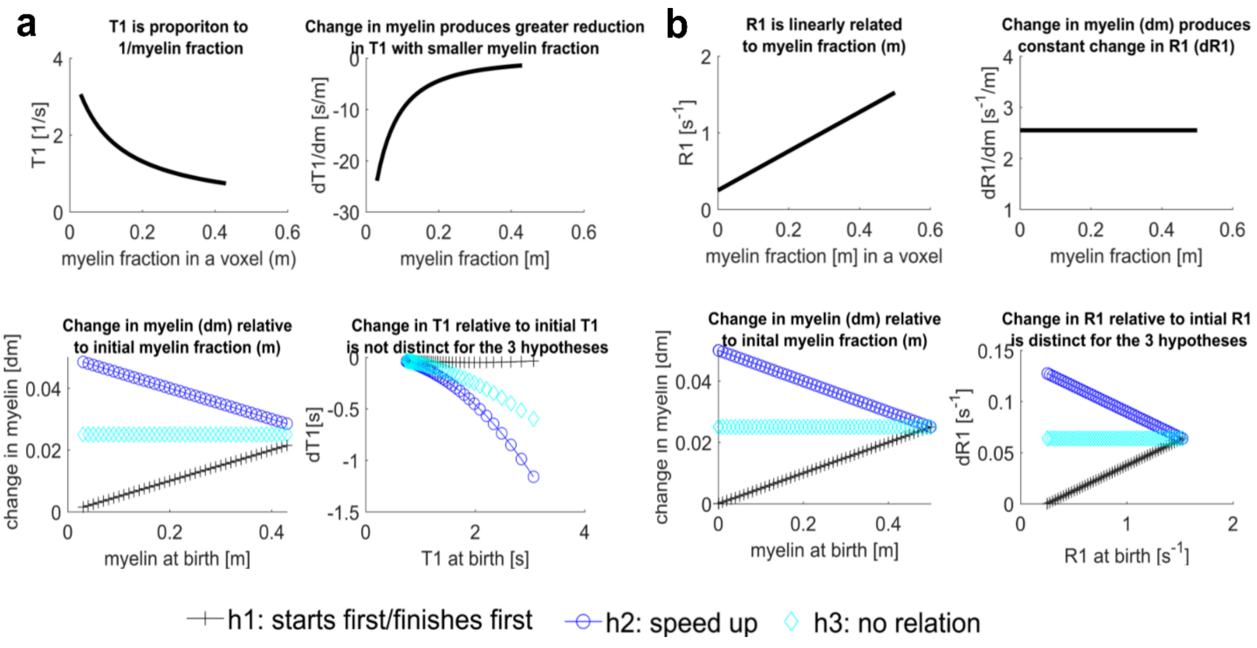


**Supplementary Figure 1.** Quantitative MRI measures of R1 (b) but not T1 (a) are linearly related to myelin content and changes in myelination over time. As such, R1 is a suitable metric to distinguish between developmental hypotheses.

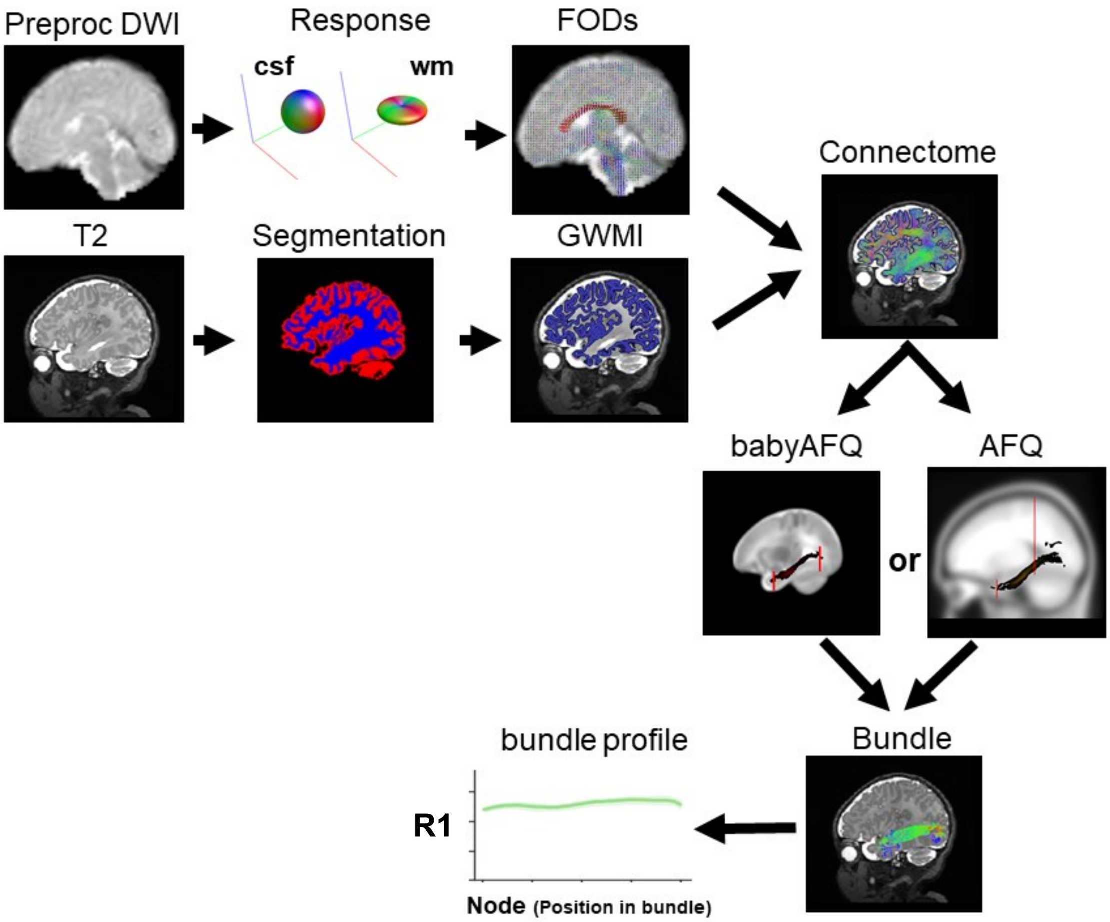


**Supplementary Figure 2.** A schematic representation of our processing pipeline. DMRI data is preprocessed and fiber density functions (FODs) are generated separately for white matter and CSF using multi-shell/multi-tissue constrained spherical deconvolution (CSD). A tissue segmentation is generated from anatomical data and used for anatomically constrained tractography (ACT). Seeds for tractography are placed at the gray/white matter interface (GWMI) and whole brain connectomes with 2 million streamlines are created for each infant and timepoint. Bundles are delineated from the whole brain connectome using either babyAFQ, which was developed here and optimized for infant data or AFQ, which was developed from adult data and serves as a benchmark. R1 development is then evaluated across the length of each bundle identified with babyAFQ.

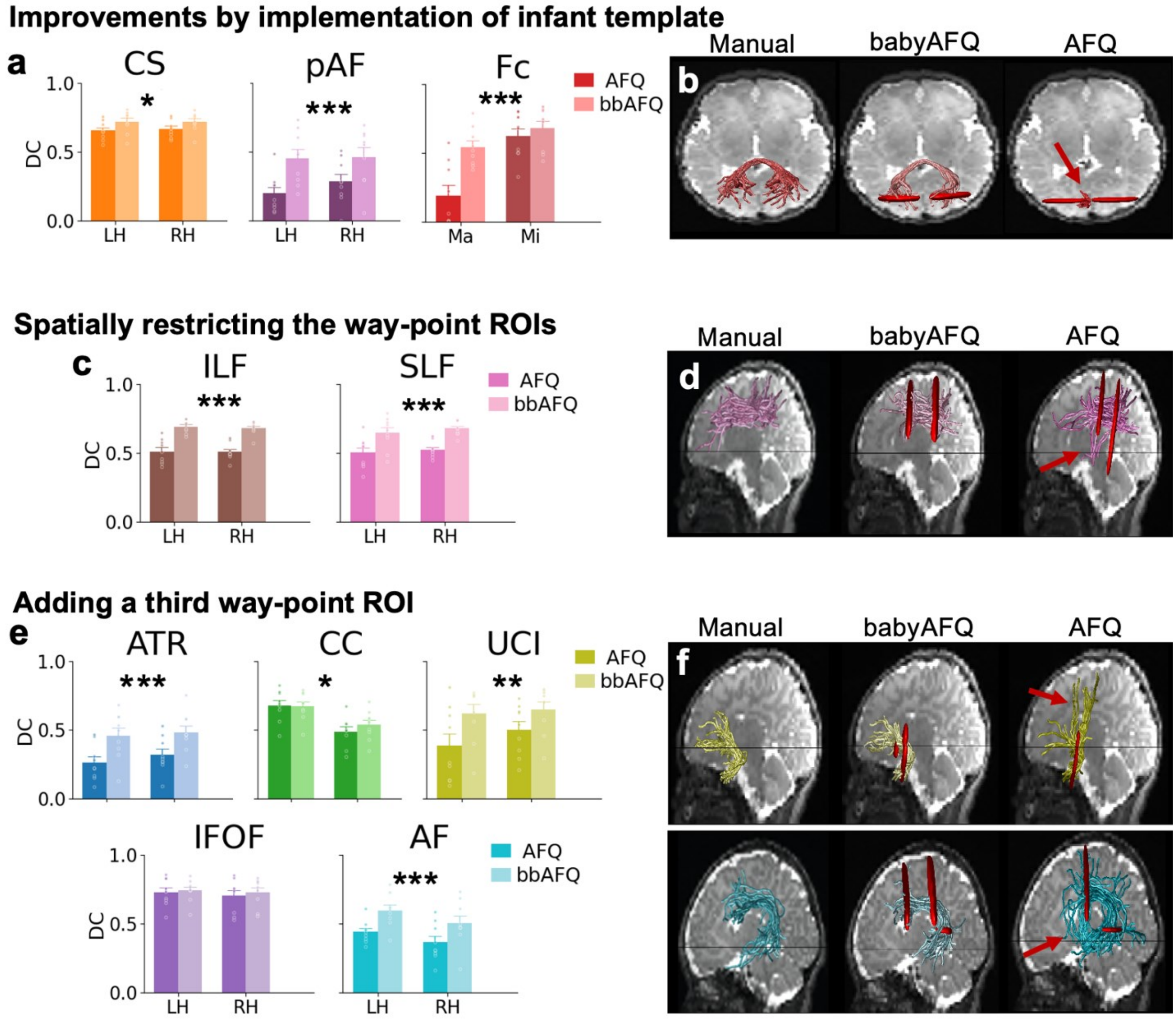


**Supplementary Figure 3.** Adapting AFQ for the infant brain (babyAFQ) improves the identification of white matter bundles in newborns. Infant-specific adaptations included the usage of an infant template brain for definition of way-point ROIs for all tracts, spatially restricted way-point ROIs for the ILF and SLF and the introduction of a third way-point ROIs for "curvy" bundles. a,c,e: Bar graphs compare AFQ and babyAFQ to manually delineated "gold-standard" bundles. Higher dice coefficients (DCs) indicate more spatial overlap with manual tracts (rmANOVAs). *$p<0.05$, **$p<0.01$, ***$p<0.001$. b,d,f: Examples of bundles delineated manually as well as with AFQ and babyAFQ presented in individual subject's left hemisphere. Abbreviations: LH: left hemisphere, RH: right hemisphere, CS: cortico-spinal tract, pAF: posterior arcuate fasciculus, Fc: forceps (Ma=forceps major; Mi=forceps minor), ILF: inferior longitudinal fasciculus, SLF: superior longitudinal fasciculus, ATR: anterior thalamic radiation, CC: cingulum cingulate, UCI: uncinate fasciculus, IFOF: inferior frontal occipital fasciculus, AF: arcuate fasciculus.

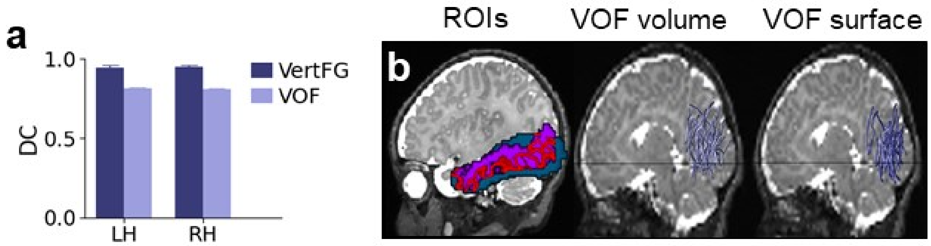


**Supplementary Figure 4.** Tracts identified as being part of the VOF are similar regardless of whether surface or volume waypoint ROIs are used. While AFQ uses surface ROIs to define the VOF, babyAFQ offers both surface (b,ROIs, purple outline) or volume (b,ROIs, blue box, gray-white matter interface in ROI in red outline) ROIs to define this bundle, as to date it is difficult to generate cortical surfaces in infants. Tracts identified as being vertical (a, dark blue) and part of the VOF (a, light blue) are highly overlapping and appear similar (b, blue) between the two types of ROIs. Abbreviations: LH: left hemisphere, RH: right hemisphere, VertFG: vertical fiber group, VOF: vertical occipital fasciculus.

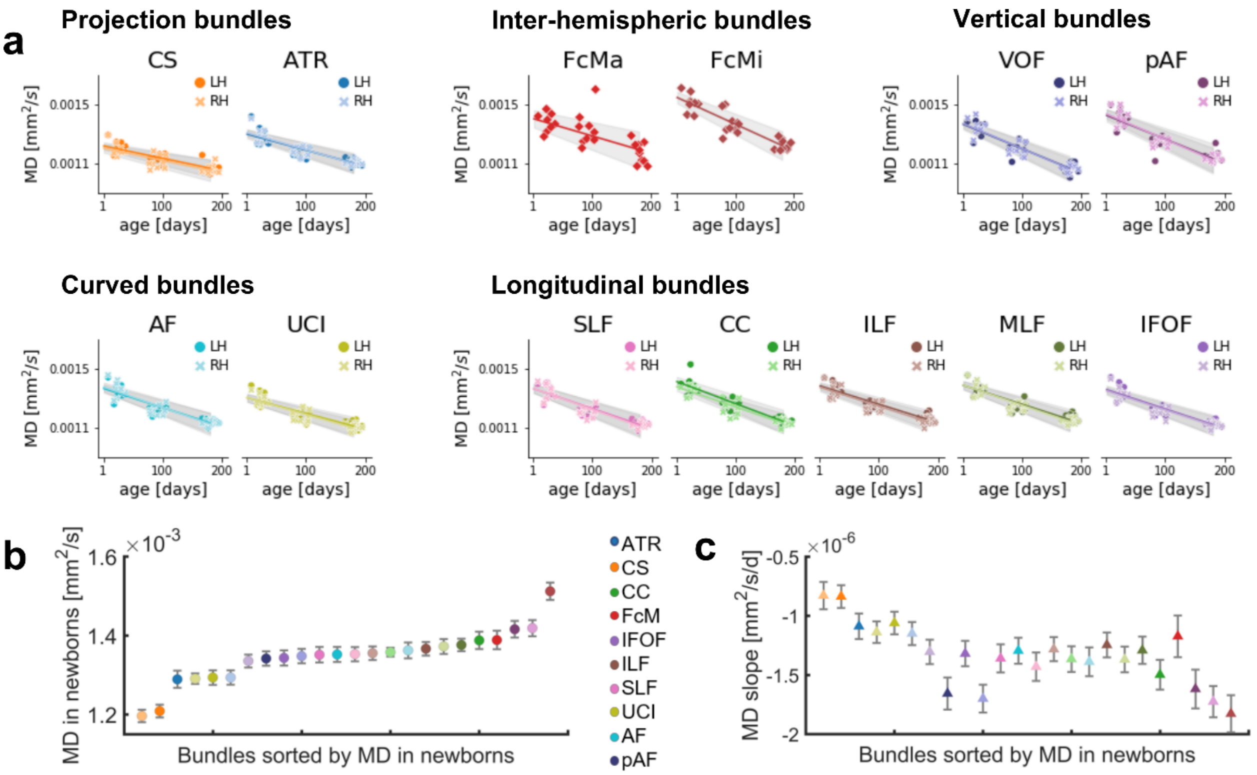


**Supplementary Figure 5.** MD of white matter bundles linearly decreases from birth to 6 months of age. a. Mean MD of each bundle as a function of age in days. Each point is a participant; markers indicate hemisphere; lines indicate LMM prediction; lines for both hemispheres fall on top of each other; gray shaded regions indicate 95% confidence intervals. b. MD measured in newborns (circles) varies across white matter bundles. c. MD development rate (slopes from LMMs, triangles) also varies across white matter bundles. Abbreviations: MD: mean diffusivity, LH: left hemisphere, RH: right hemisphere, CS: cortico-spinal tract, ATR: anterior thalamic radiation, FcMa: forceps major; FcMi: forceps minor, VOF: vertical occipital fasciculus, pAF: posterior arcuate fasciculus, AF: arcuate fasciculus, UCI: uncinate fasciculus, SLF: superior longitudinal fasciculus, CC: cingulum cingulate, ILF: inferior longitudinal fasciculus, MLF: middle longitudinal fasciculus, IFOF: inferior frontal occipital fasciculus.

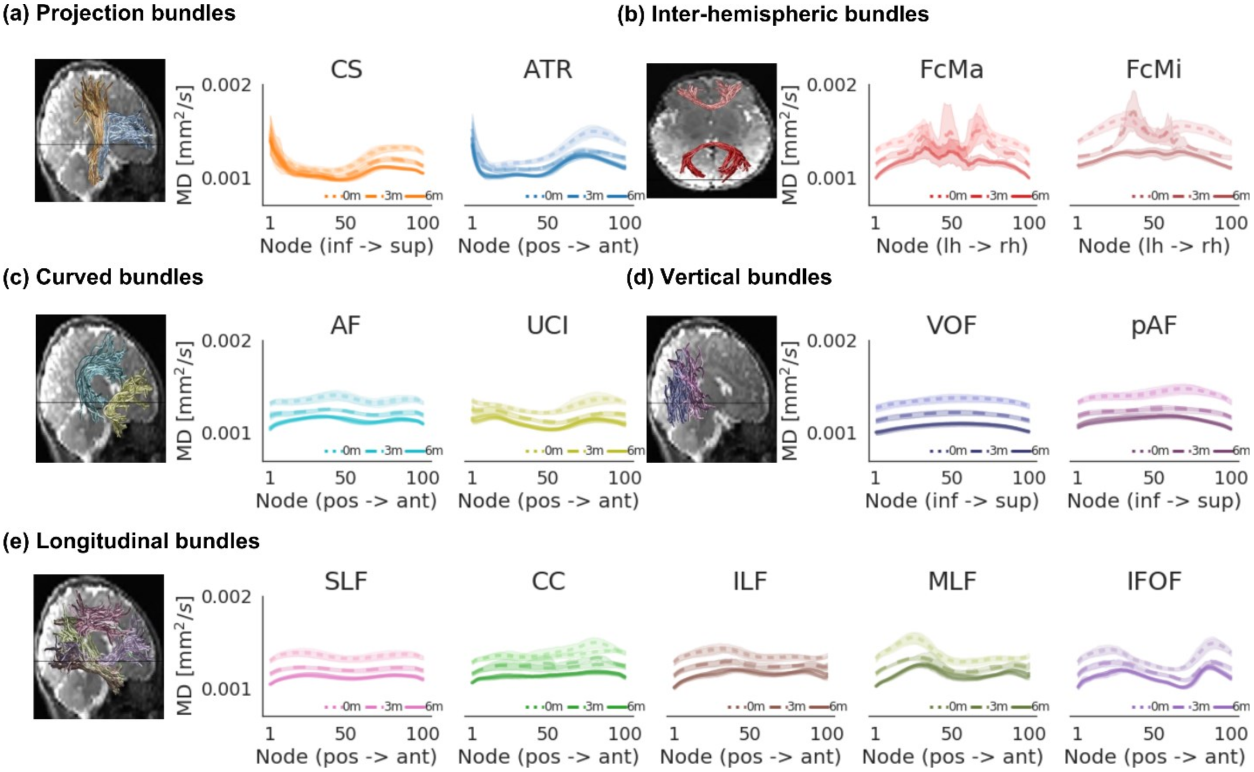


**Supplementary Figure 6.** Development of MD along each bundle. Mean MD across infants is displayed in both hemispheres (lines for the two hemispheres fall on top of each other) along the length of each bundle in newborns (0m, dotted line), 3-months-olds (3m, dashed line), and 6-months-olds (6m, solid line). Shaded regions: 95% confidence intervals. Left panels show the bundles in a representative newborn. Abbreviations: MD: mean diffusivity, CS: cortico-spinal tract, ATR: anterior thalamic radiation, FcMa: forceps major; FcMi: forceps minor, VOF: vertical occipital fasciculus, pAF: posterior arcuate fasciculus, AF: arcuate fasciculus, UCI: uncinate fasciculus, SLF: superior longitudinal fasciculus, CC: cingulum cingulate, ILF: inferior longitudinal fasciculus, MLF: middle longitudinal fasciculus, IFOF: inferior frontal occipital fasciculus.

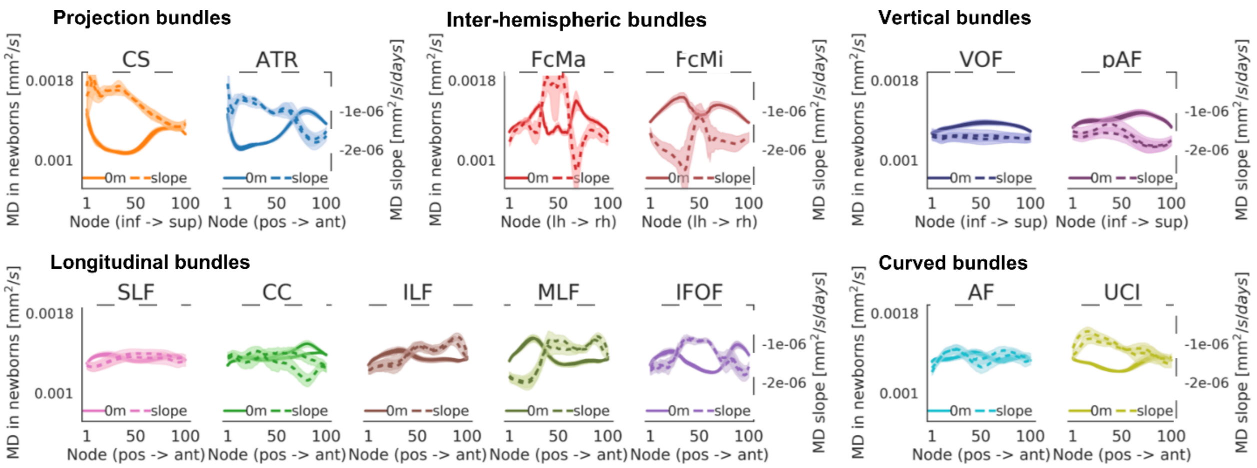


**Supplementary Figure 7.** MD development rate varies along the length of each bundle. a. Each panel jointly shows measured MD in newborns (left y axis, solid line) and the slope of MD development (right y axis, dashed line) at each node along the bundle. Faster development (more negative slope) corresponds to lower values of dashed lines. Higher MD in newborns correspond to higher values in solid lines. Lines from both hemispheres are presented separately but fall on top of each other. Abbreviations: MD: mean diffusivity, CS: cortico-spinal tract, ATR: anterior thalamic radiation, FcMa: forceps major; FcMi: forceps minor, VOF: vertical occipital fasciculus, pAF: posterior arcuate fasciculus, AF: arcuate fasciculus, UCI: uncinate fasciculus, SLF: superior longitudinal fasciculus, CC: cingulum cingulate, ILF: inferior longitudinal fasciculus, MLF: middle longitudinal fasciculus, IFOF: inferior frontal occipital fasciculus.

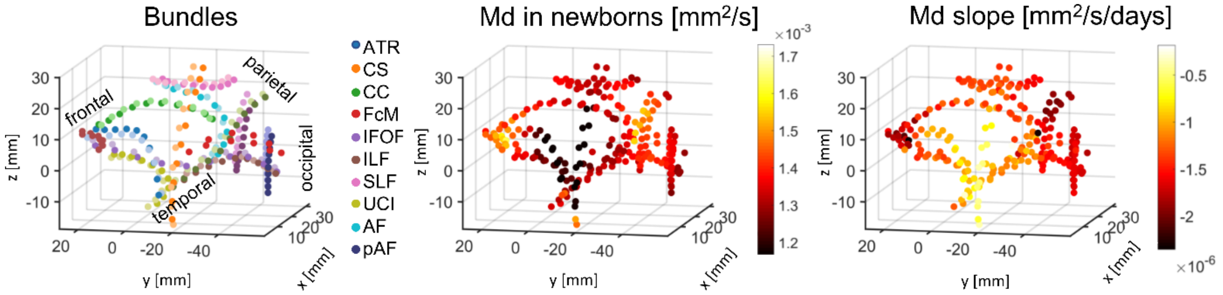


**Supplementary Figure 8.** Spatial gradients and MD at birth together explain MD development. Left: A 3D plot of the spatial layout of every 10th node of all bundles. Nodes are color coded by bundle membership and approximate lobe labels are included to provide orientation. Middle: A 3D plot of the spatial layout of MD in newborns. Right: A 3D plot of the spatial layout of slopes of MD development. Joined evaluation of the subplots suggests that MD slopes vary in space, and are related to MD in newborns. We quantitatively tested these observations using an LMM relating MD slope at every 10th node to measured MD in newborns and spatial coordinate: |x|, y, z, |x|*y, |x|*z, and z*y (analysis was done across all nodes, with a random intercept per bundle). This combined model showed a significant negative relationship between the rate of MD development and MD in newborns: ($\beta$=-0.002; p<0.0001) and significant effects of spatial location along the x-axis ($\beta$=-9.58x$10^{-8}$, p=0.0004), the y-axis ($\beta$=9.78x$10^{-8}$, p<0.0001), the z-axis ($\beta$=-1.56x$10^{-7}$, p<0.0001), and the |x|*y axis ($\beta$=6.41*$10^{-8}$, p=0.03). Overall, this combined model explains 71% of the variance of the rate of MD development ($R^2$=0.71). Abbreviations: MD: mean diffusivity, CS: cortico-spinal tract, ATR: anterior thalamic radiation, FcMa: forceps major; FcMi: forceps minor, VOF: vertical occipital fasciculus, pAF: posterior arcuate fasciculus, AF: arcuate fasciculus, UCI: uncinate fasciculus, SLF: superior longitudinal fasciculus, CC: cingulum cingulate, ILF: inferior longitudinal fasciculus, MLF: middle longitudinal fasciculus, IFOF: inferior frontal occipital fasciculus.

**Supplementary Figure 9.** BabyAFQ successfully identifies all bundles in all infants and timepoints. On the following pages, we show each individual's bundles identified with babyAFQ as well as AFQ and manually (where applicable). Data is sorted by bundle. Bundles identified with babyAFQ show the expected spatial extent and trajectory throughout.

#### Left Anterior Thalamic Radiation
#### Newborn

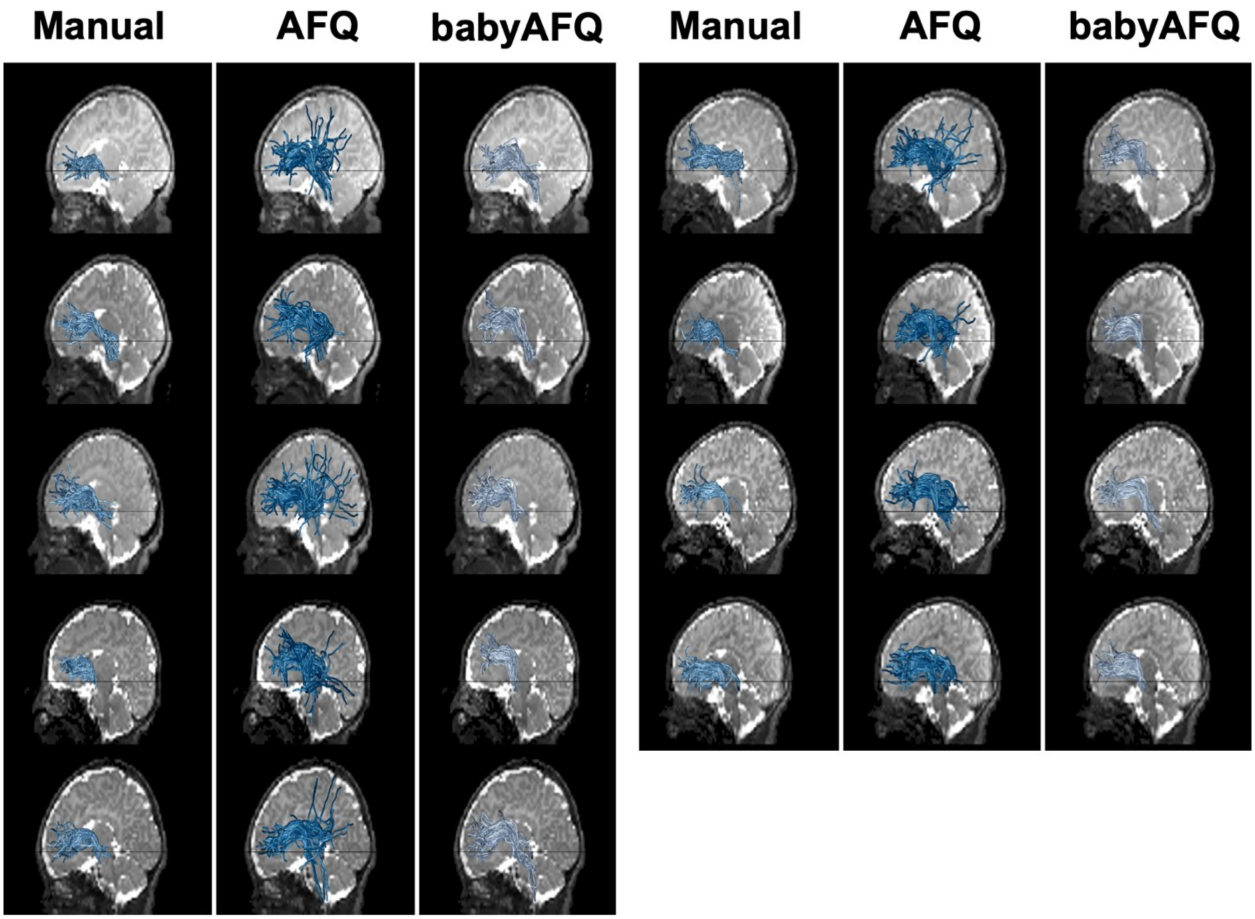

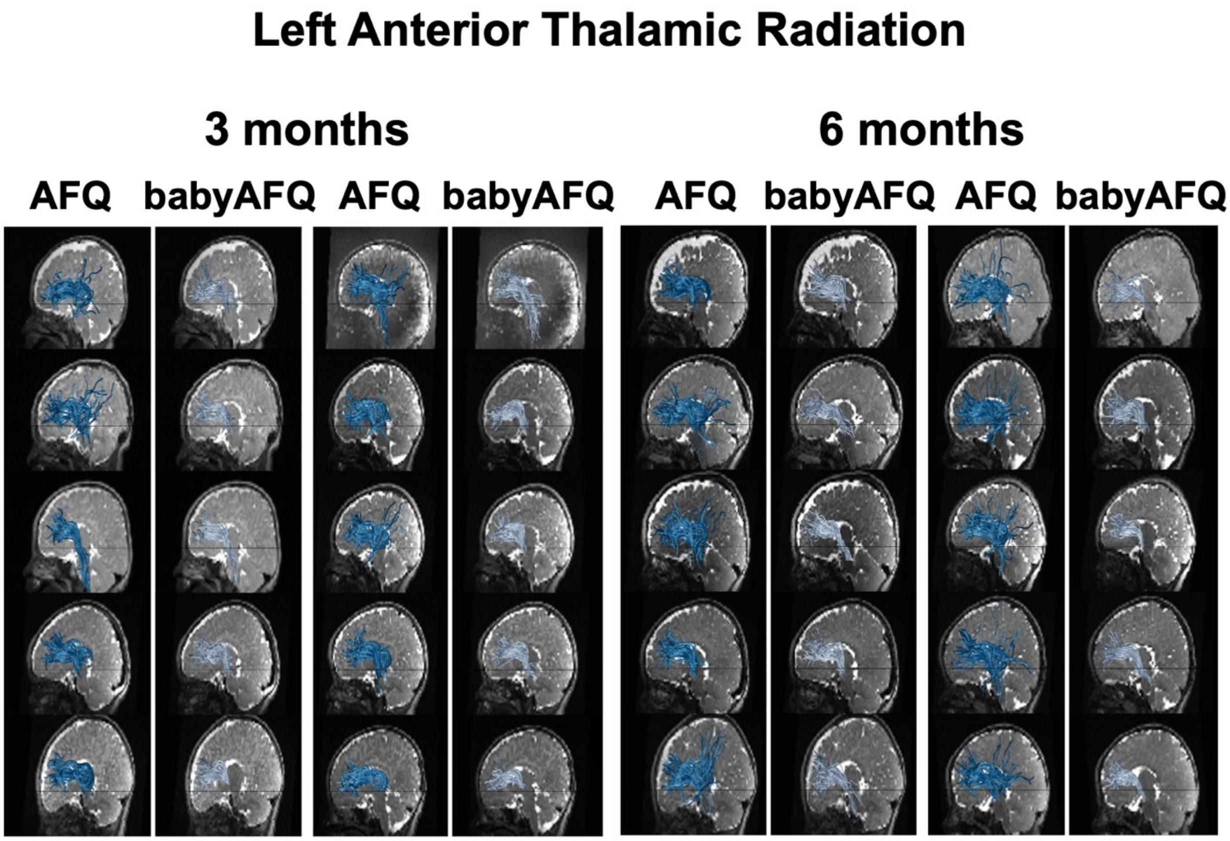
Left Anterior Thalamic Radiation
3 months
6 months
AFQ
babyAFQ
AFQ
babyAFQ
AFQ
babyAFQ
AFQ
babyAFQ

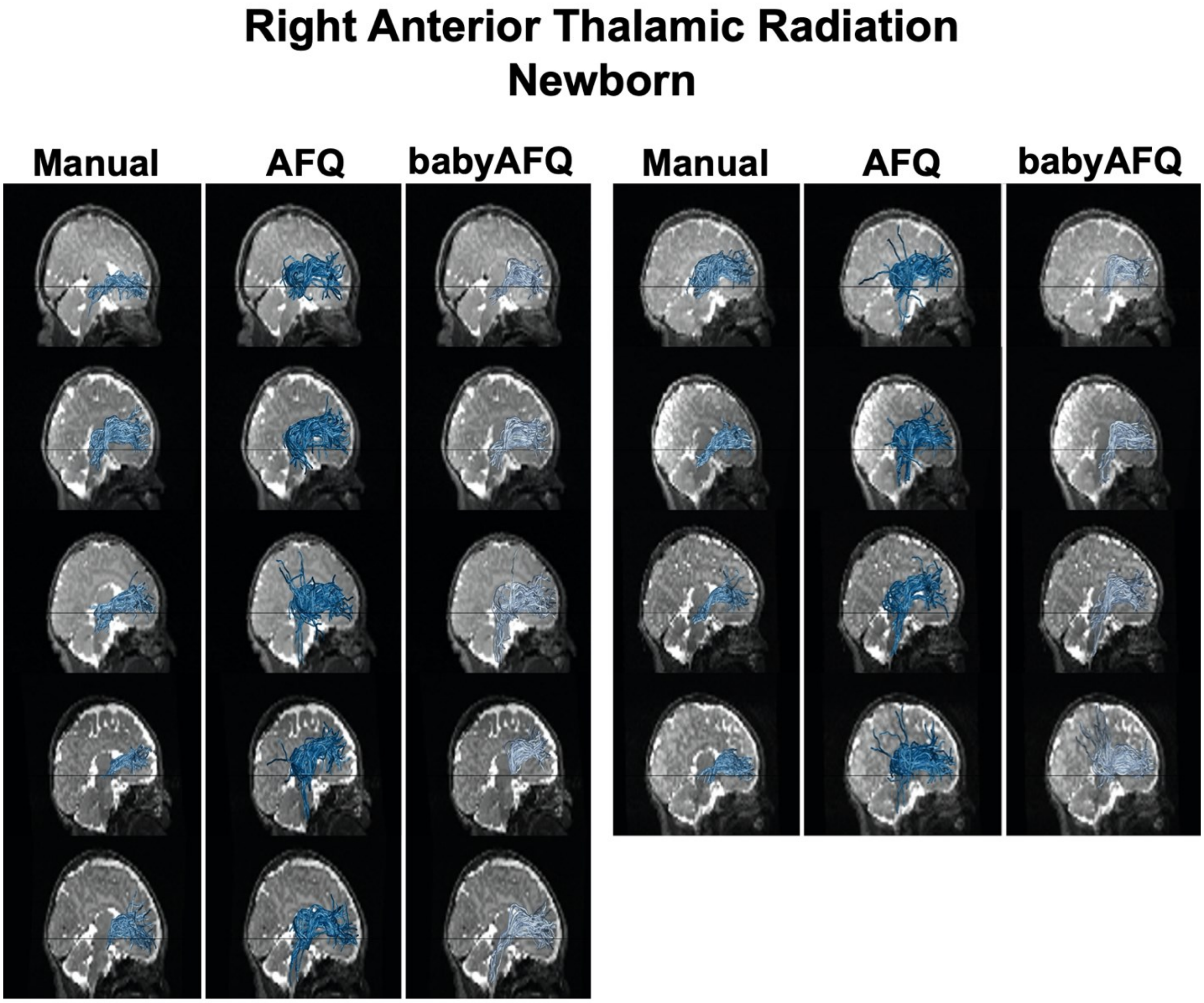
Right Anterior Thalamic Radiation
Newborn
Manual
AFQ
babyAFQ
Manual
AFQ
babyAFQ

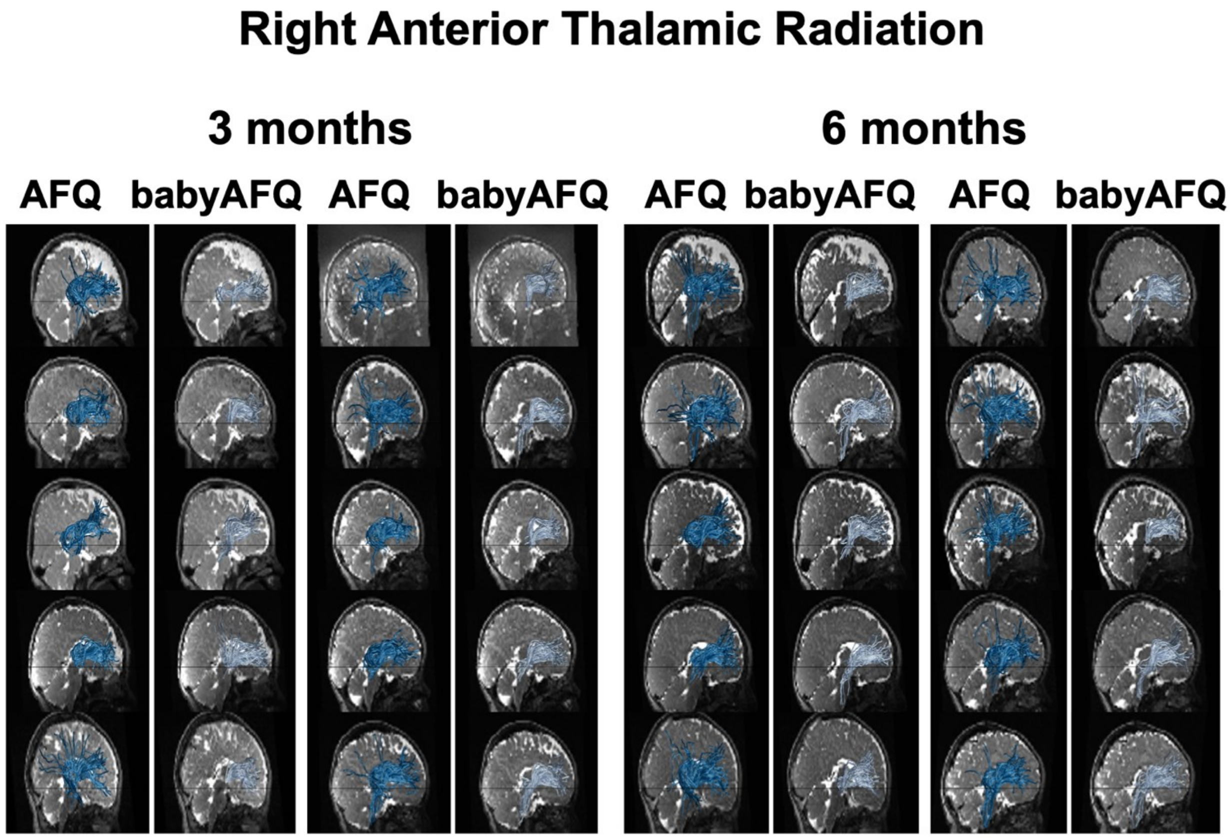
Right Anterior Thalamic Radiation
3 months
6 months
AFQ
babyAFQ
AFQ
babyAFQ
AFQ
babyAFQ
AFQ
babyAFQ

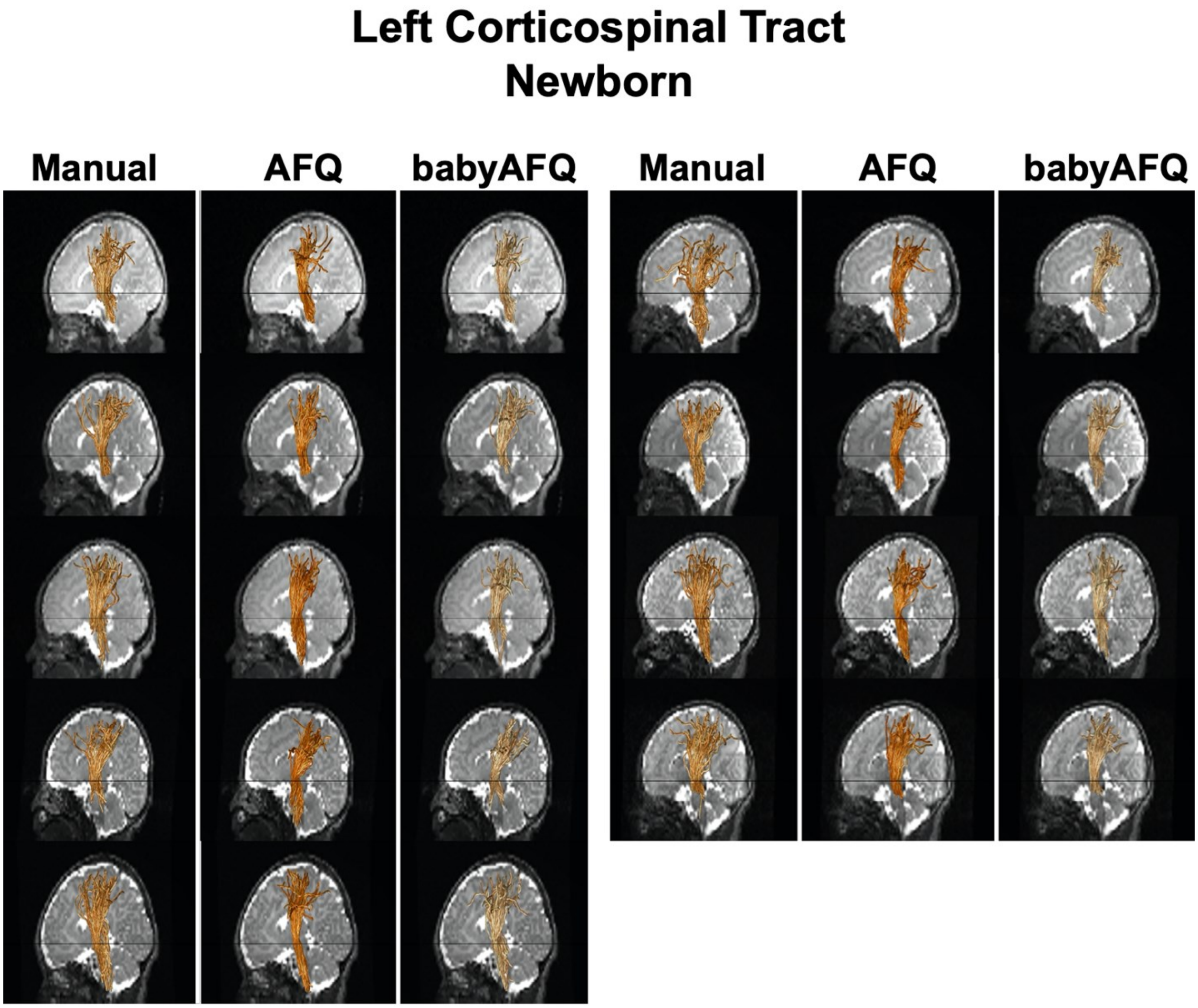
Left Corticospinal Tract
Newborn
Manual
AFQ
babyAFQ
Manual
AFQ
babyAFQ

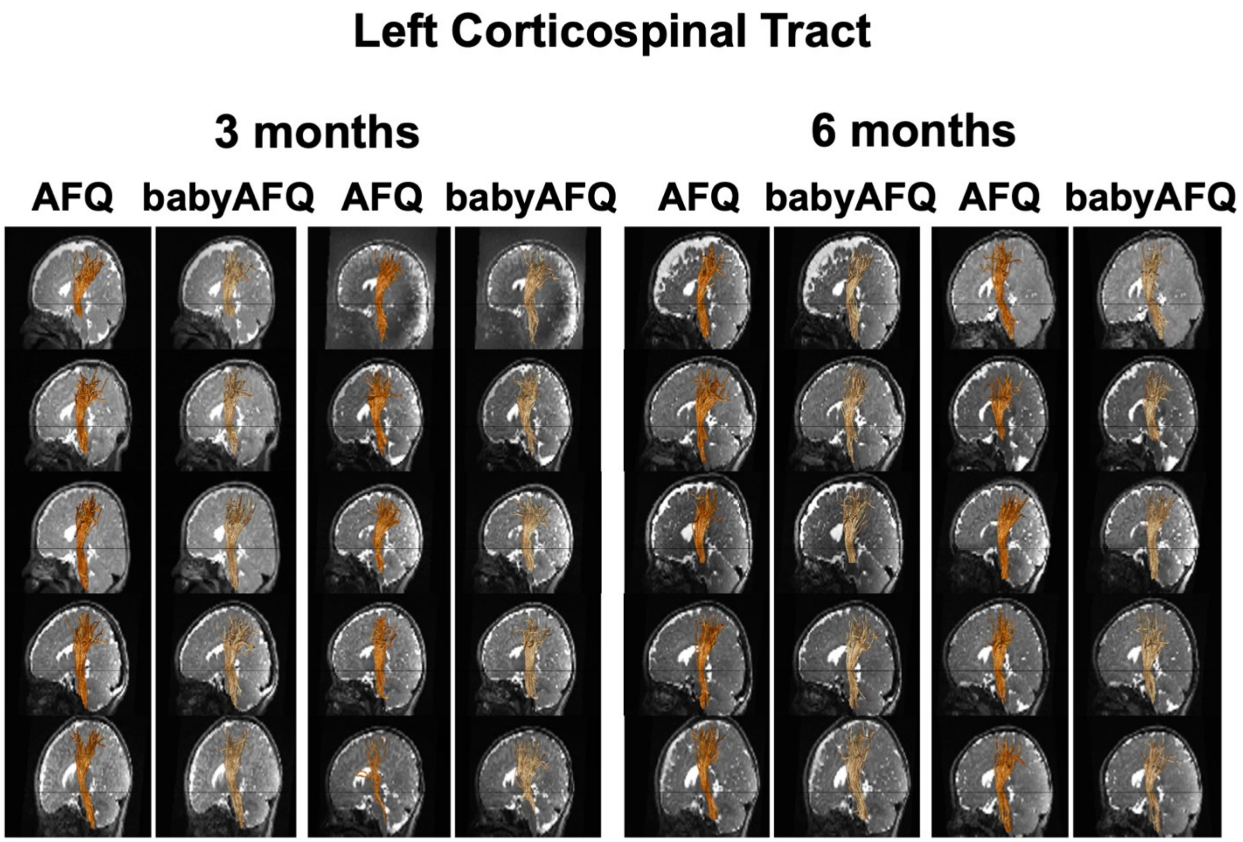
Left Corticospinal Tract
3 months
6 months
AFQ
babyAFQ
AFQ
babyAFQ
AFQ
babyAFQ
AFQ
babyAFQ

#### Right Corticospinal Tract
#### Newborn

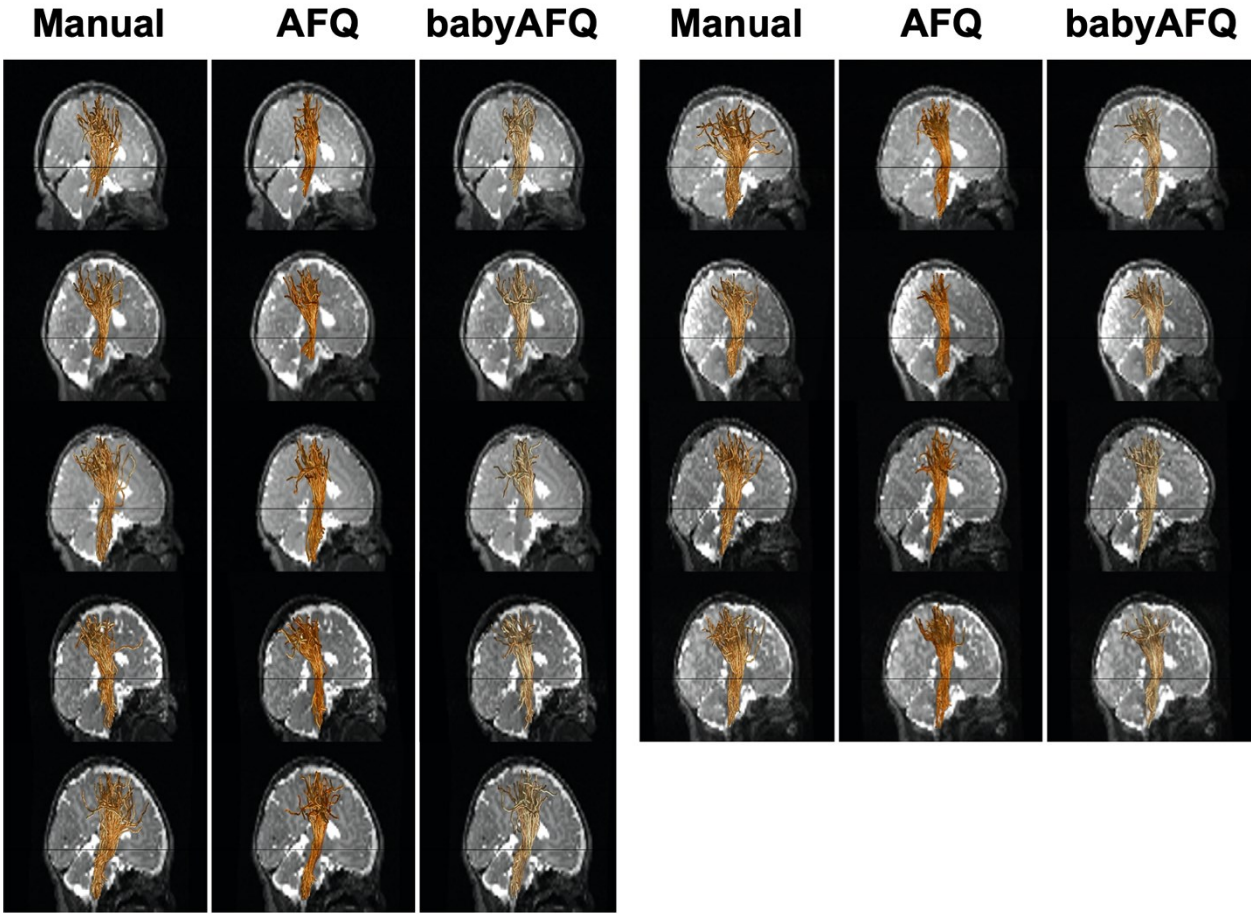

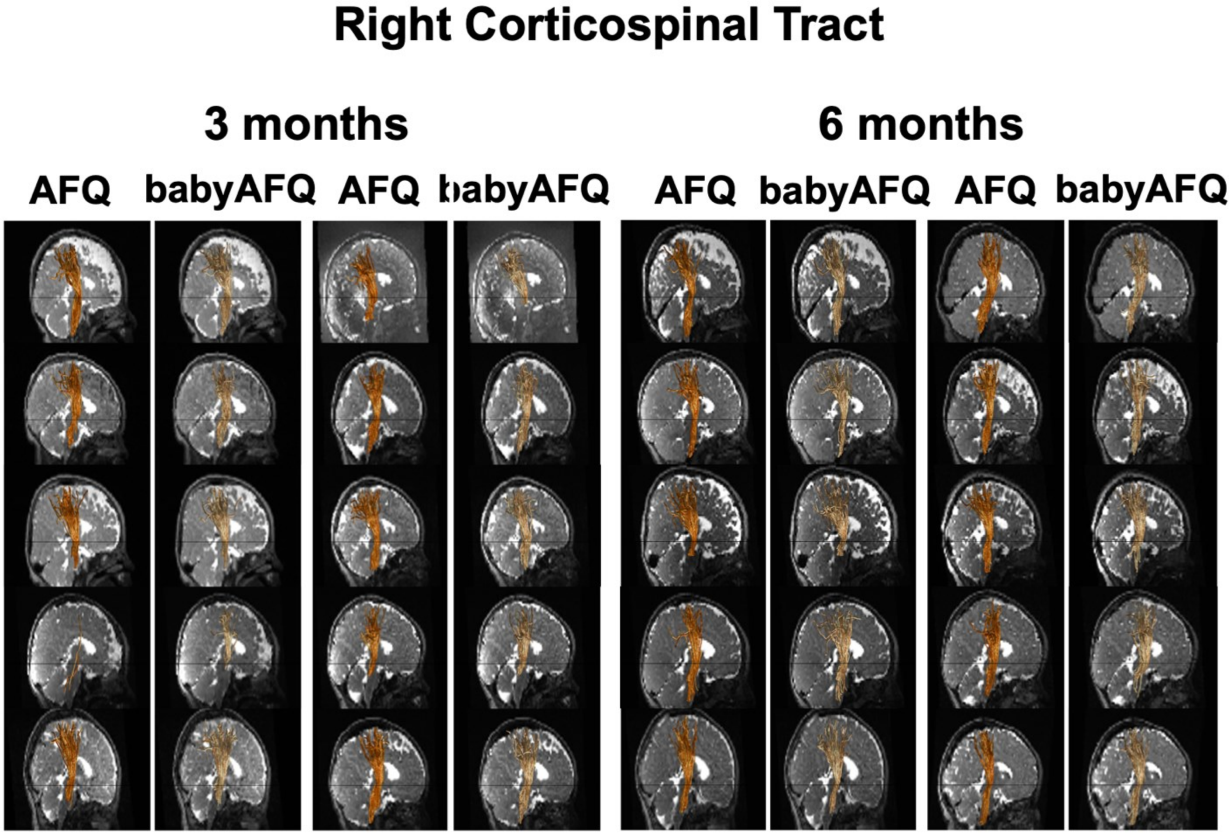

Right Corticospinal Tract
3 months
6 months
AFQ
babyAFQ
AFQ
babyAFQ
AFQ
babyAFQ
AFQ
babyAFQ

#### Left Cingulum Cingulate
#### Newborn

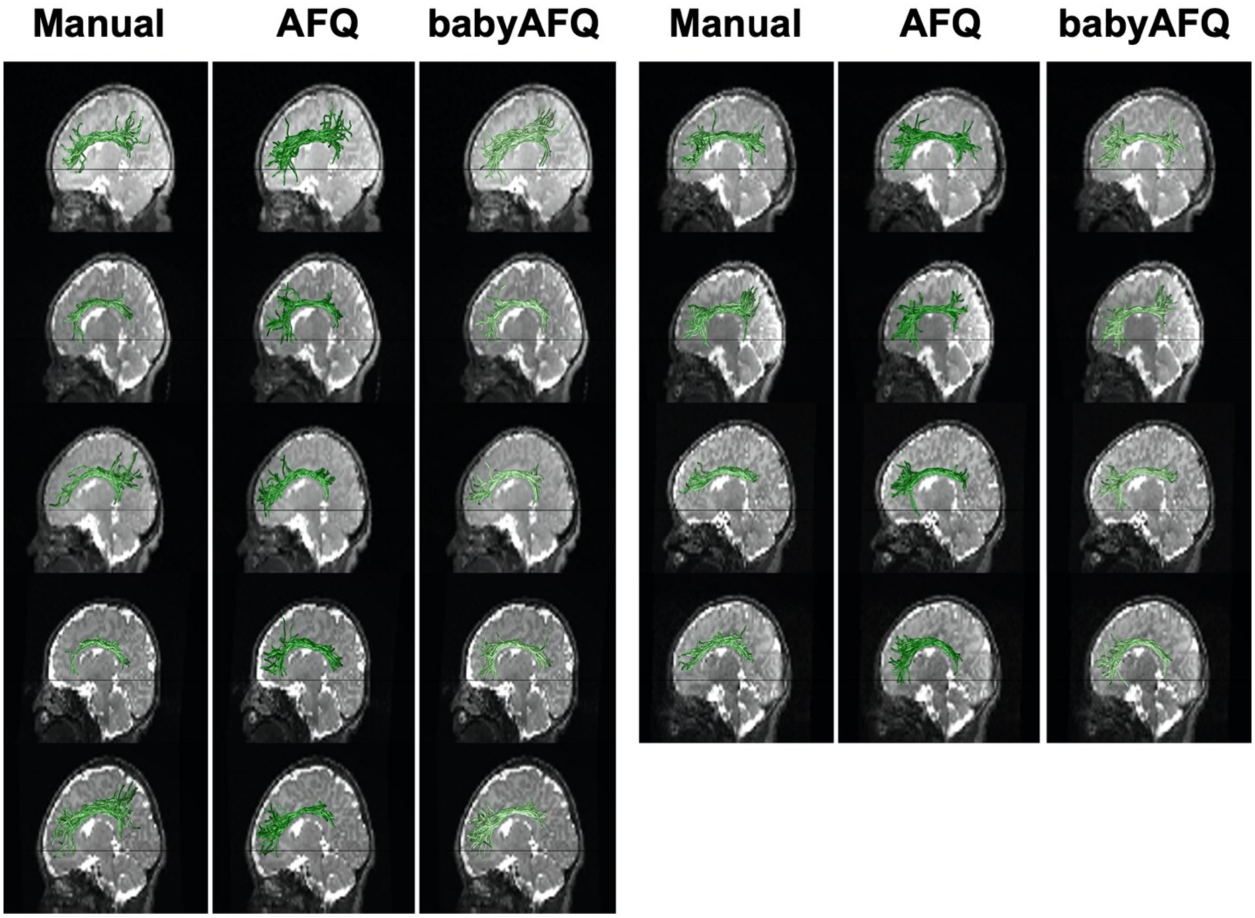

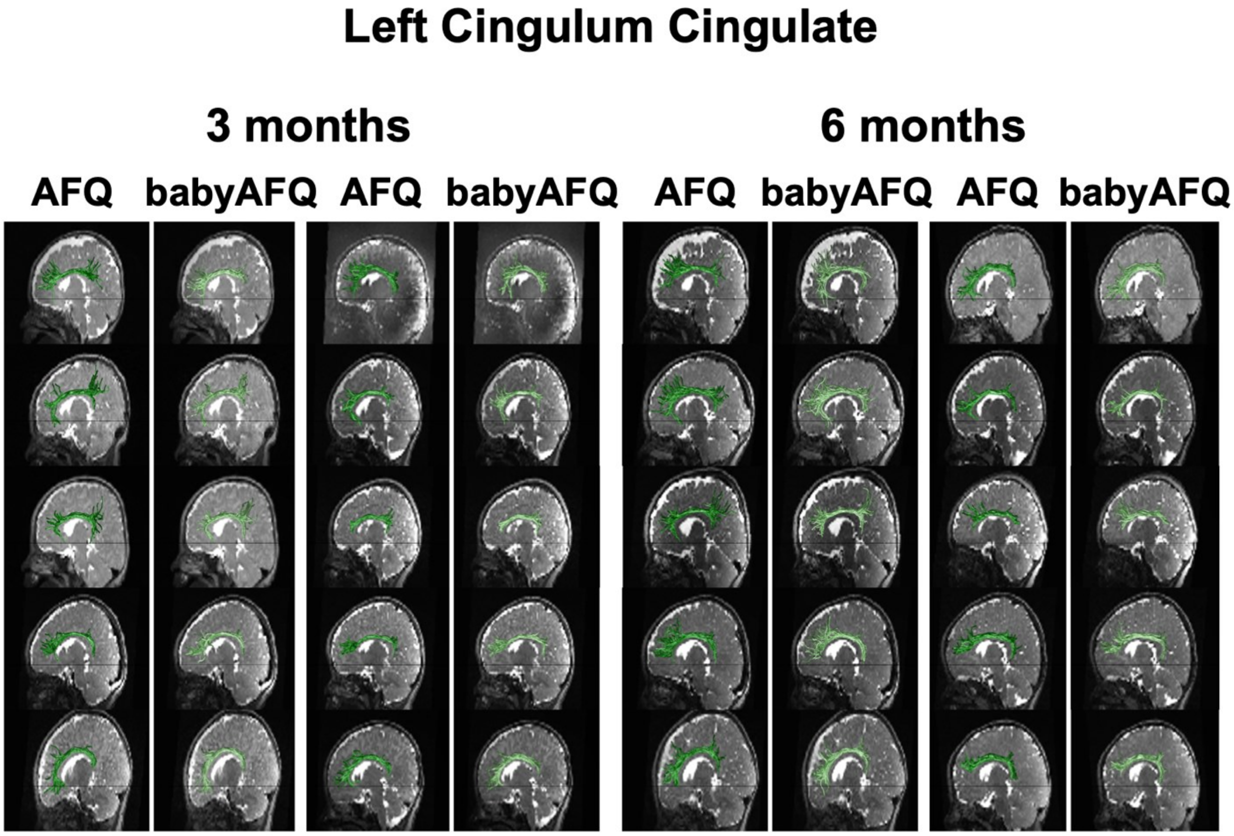
Left Cingulum Cingulate
3 months
6 months
AFQ
babyAFQ
AFQ
babyAFQ
AFQ
babyAFQ
AFQ
babyAFQ

#### Right Cingulum Cingulate
#### Newborn

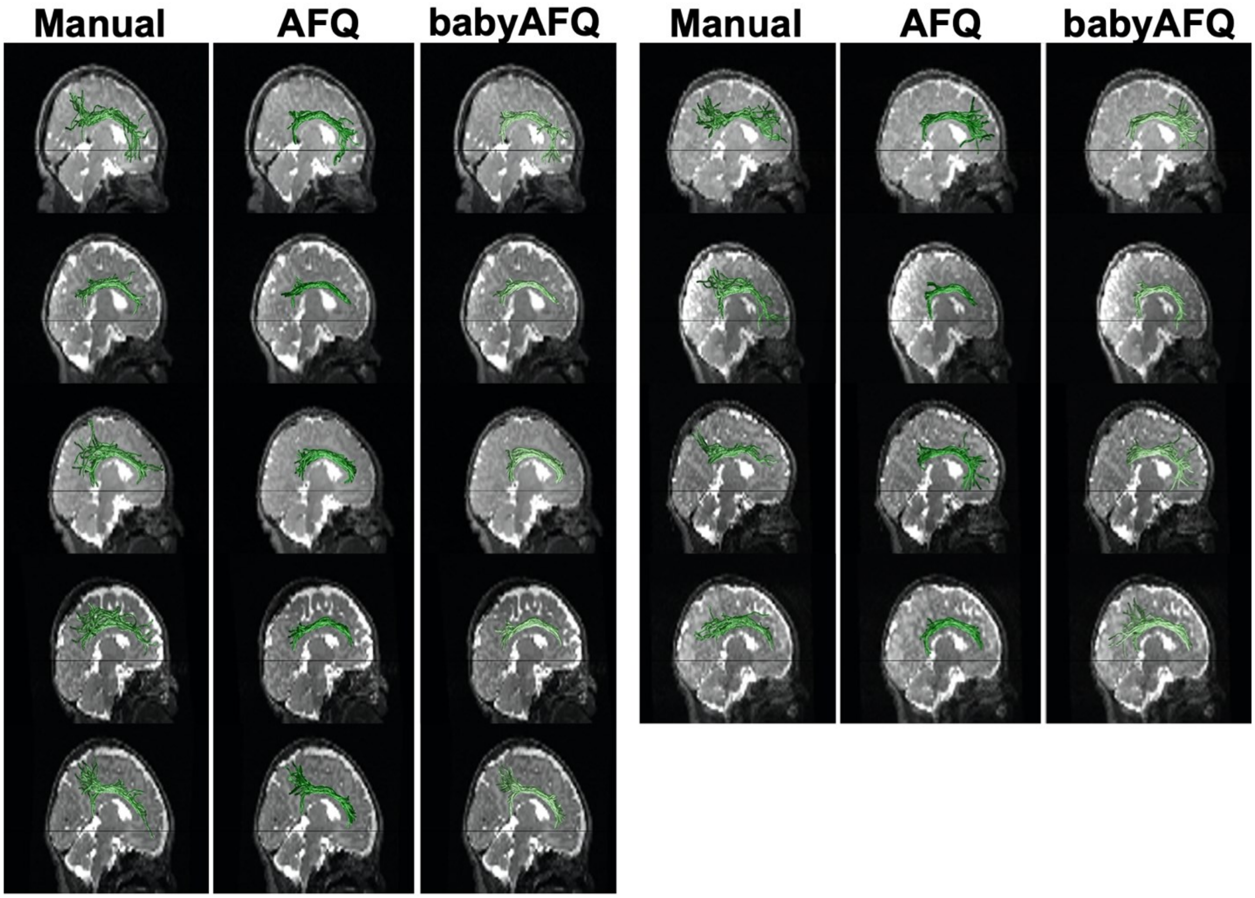

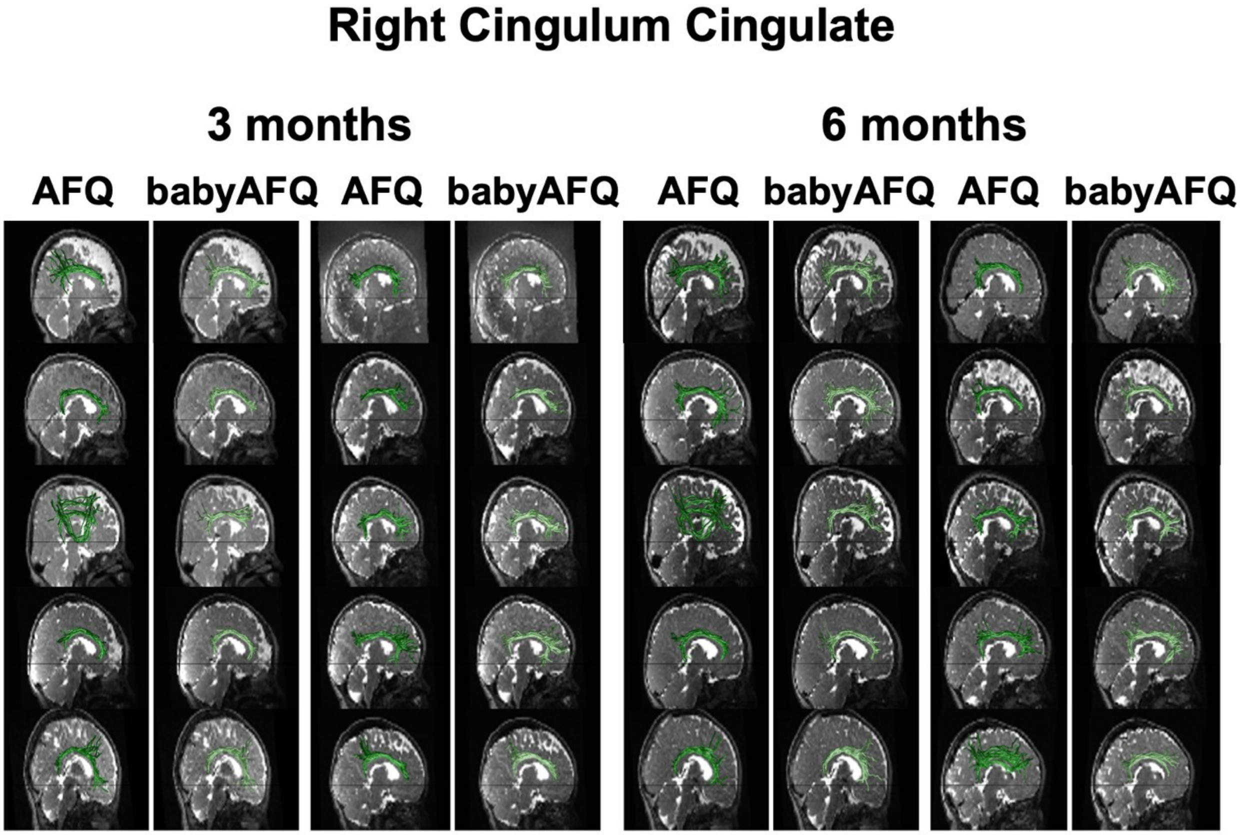
Right Cingulum Cingulate
3 months
6 months
AFQ
babyAFQ
AFQ
babyAFQ
AFQ
babyAFQ
AFQ
babyAFQ

**Forceps Major**
**Newborn**

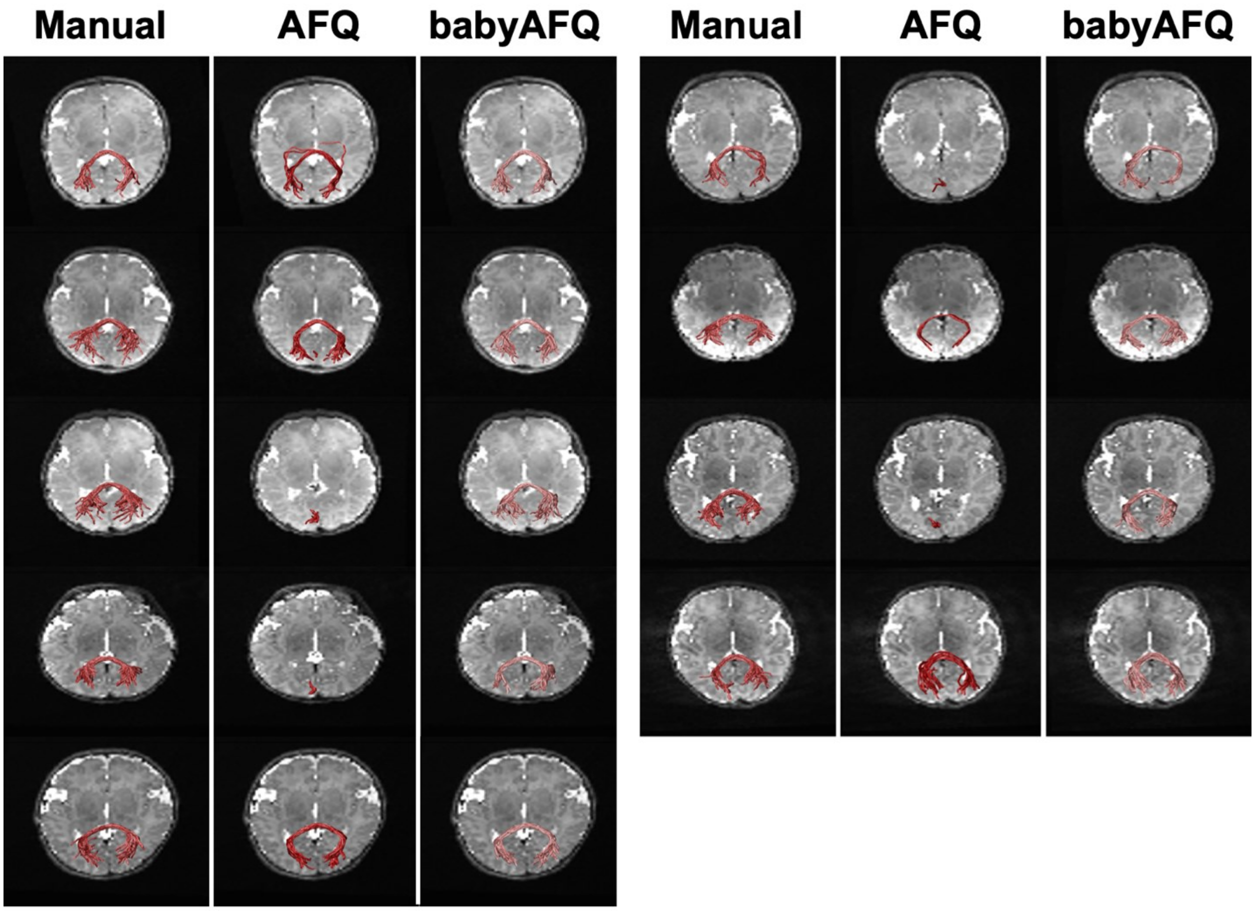

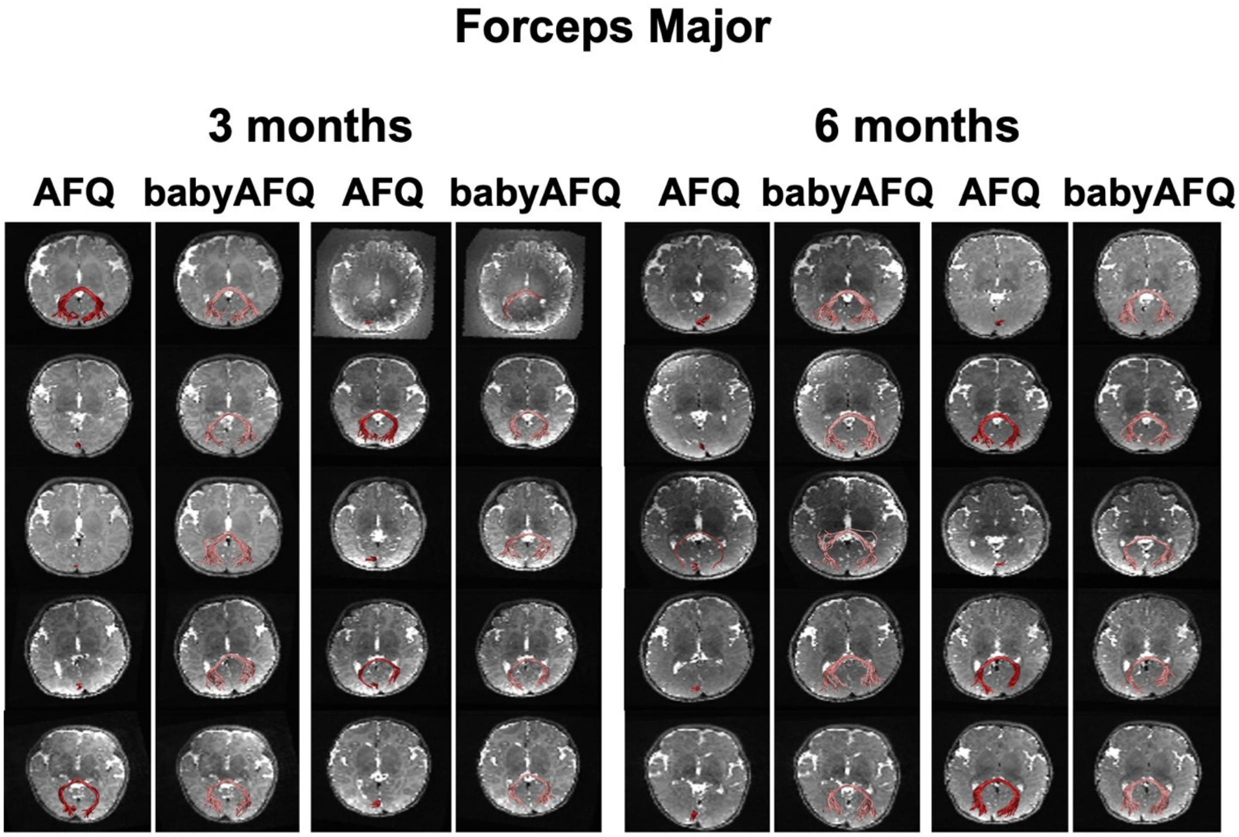
Forceps Major
3 months
6 months
AFQ
babyAFQ
AFQ
babyAFQ
AFQ
babyAFQ
AFQ
babyAFQ

**Forceps Minor**
**Newborn**

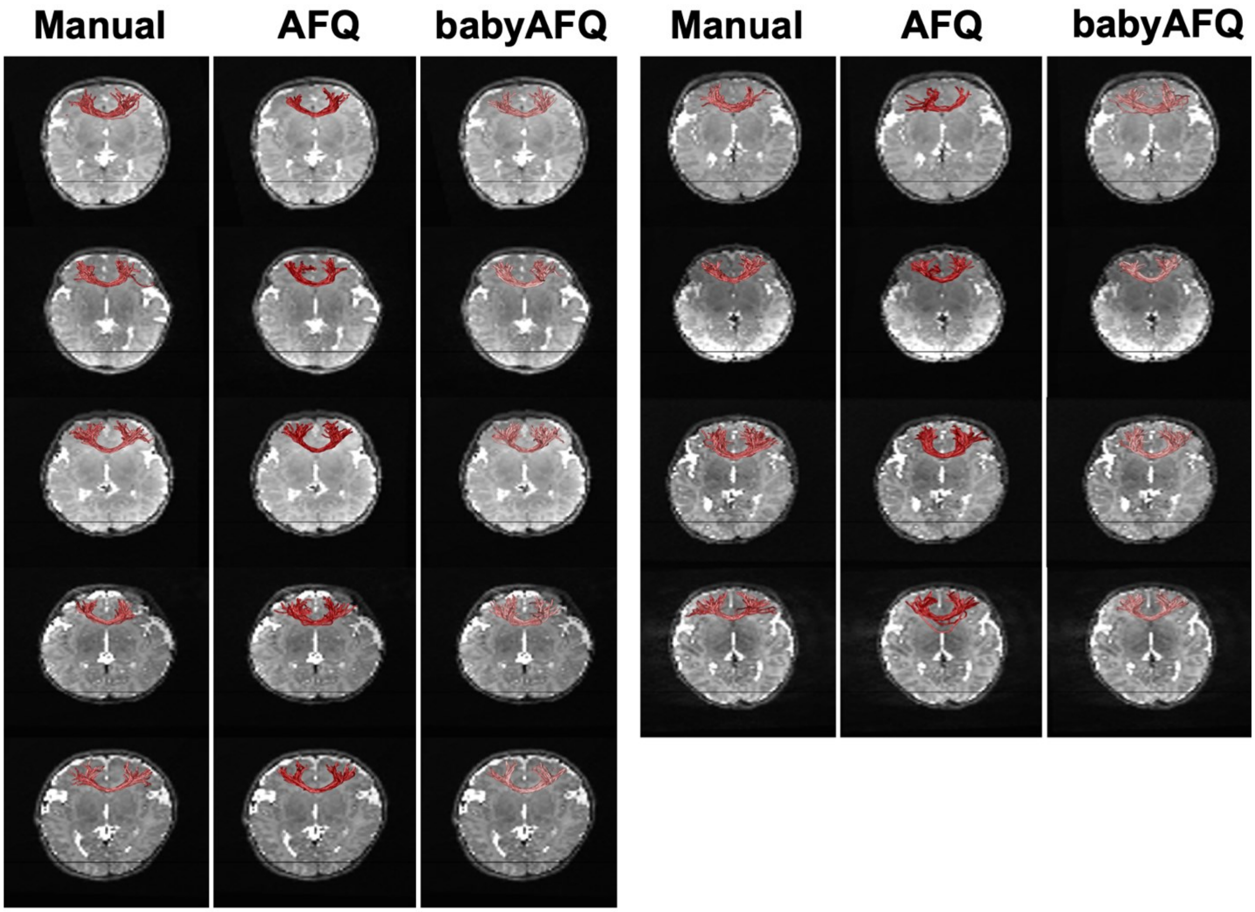

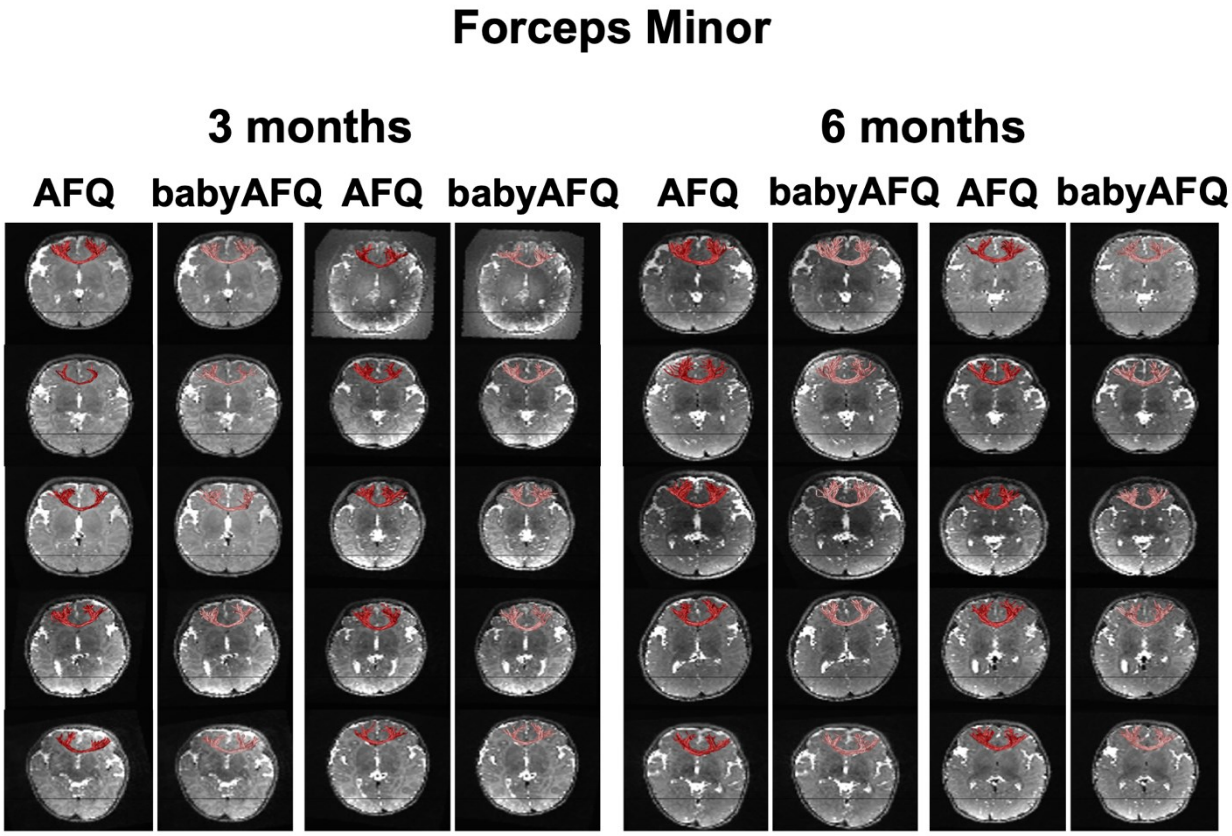
Forceps Minor
3 months
6 months
AFQ
babyAFQ
AFQ
babyAFQ
AFQ
babyAFQ
AFQ
babyAFQ

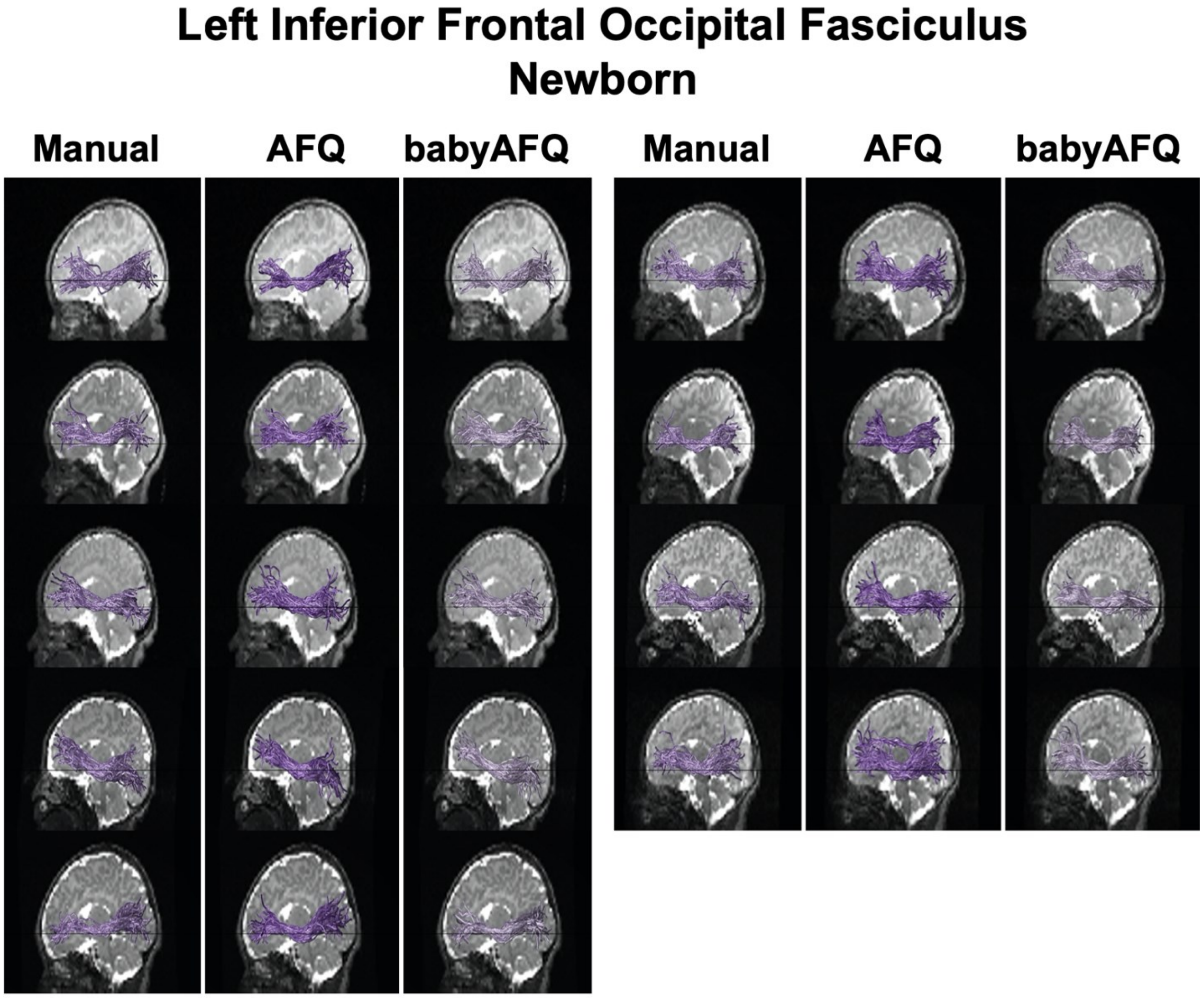
Left Inferior Frontal Occipital Fasciculus
Newborn
Manual
AFQ
babyAFQ
Manual
AFQ
babyAFQ

#### Left Inferior Frontal Occipital Fasciculus

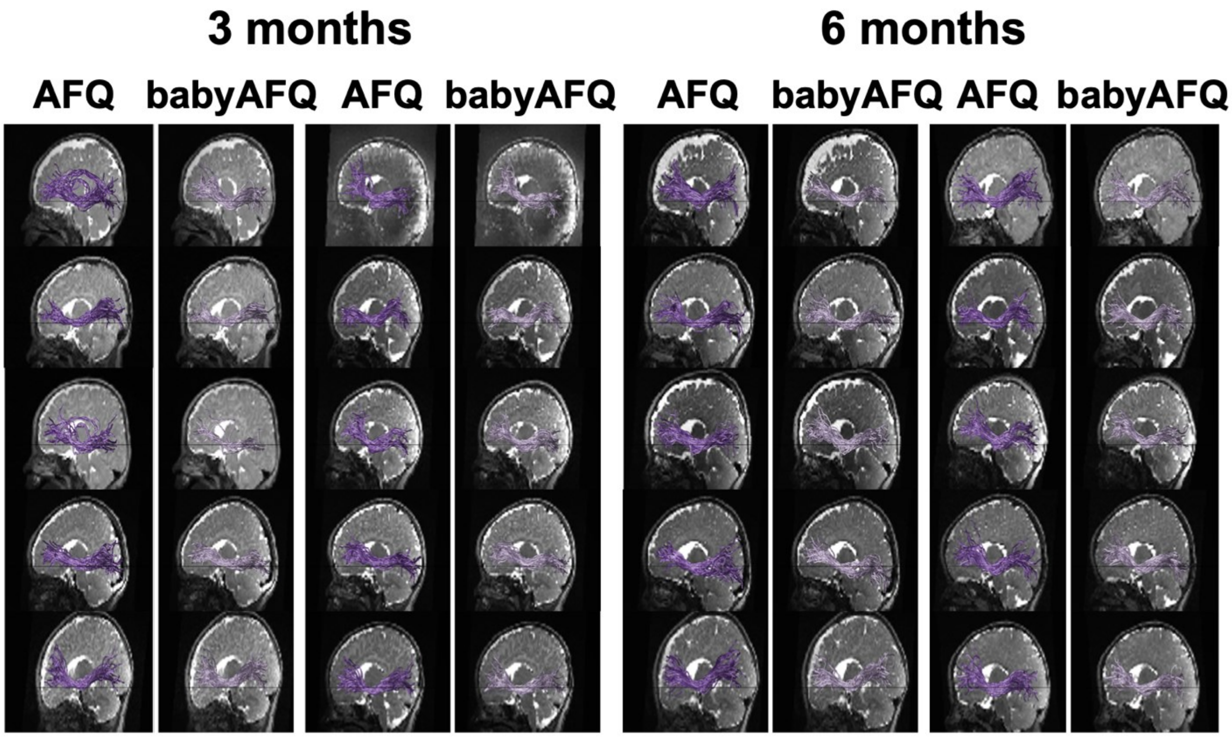

#### Right Inferior Frontal Occipital Fasciculus
#### Newborn

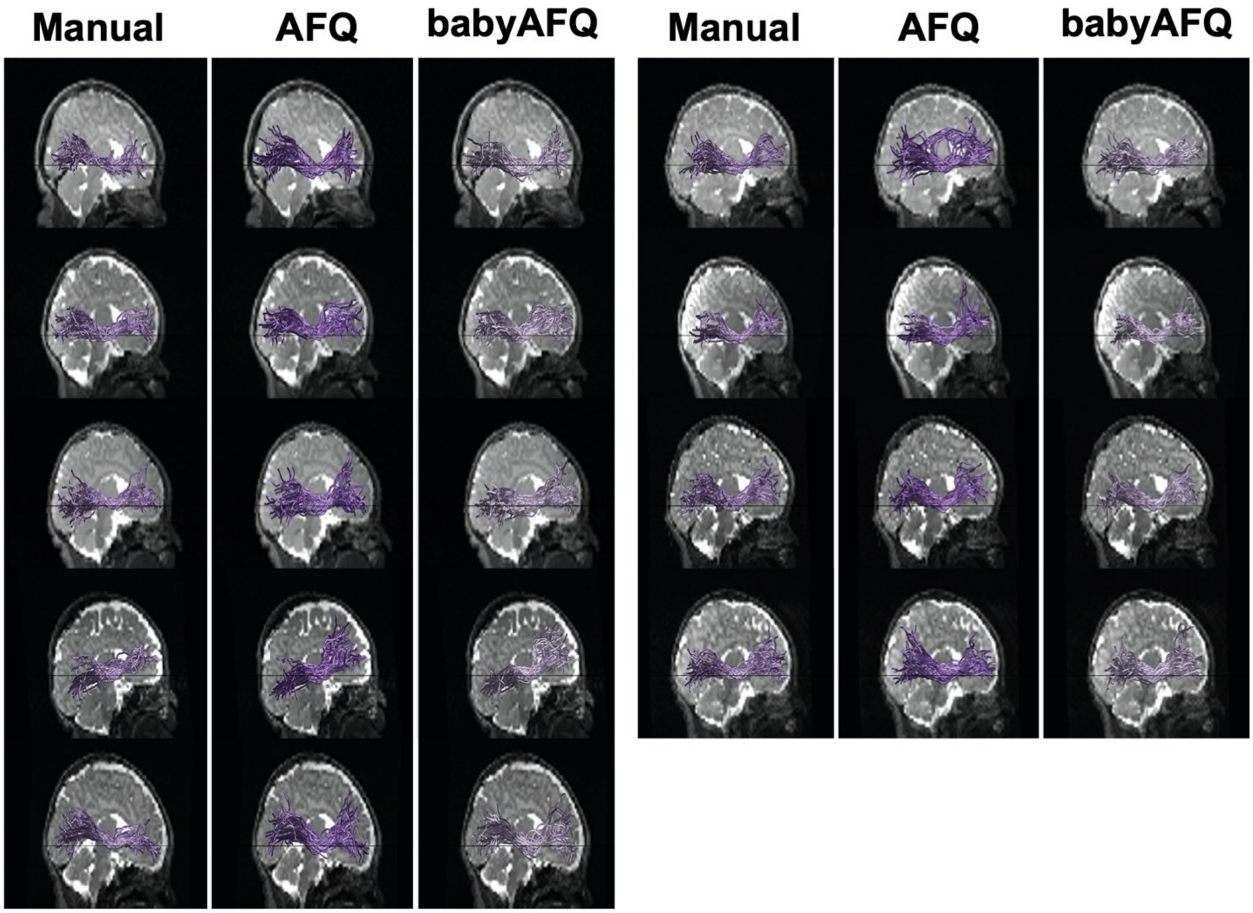

#### Right Inferior Frontal Occipital Fasciculus

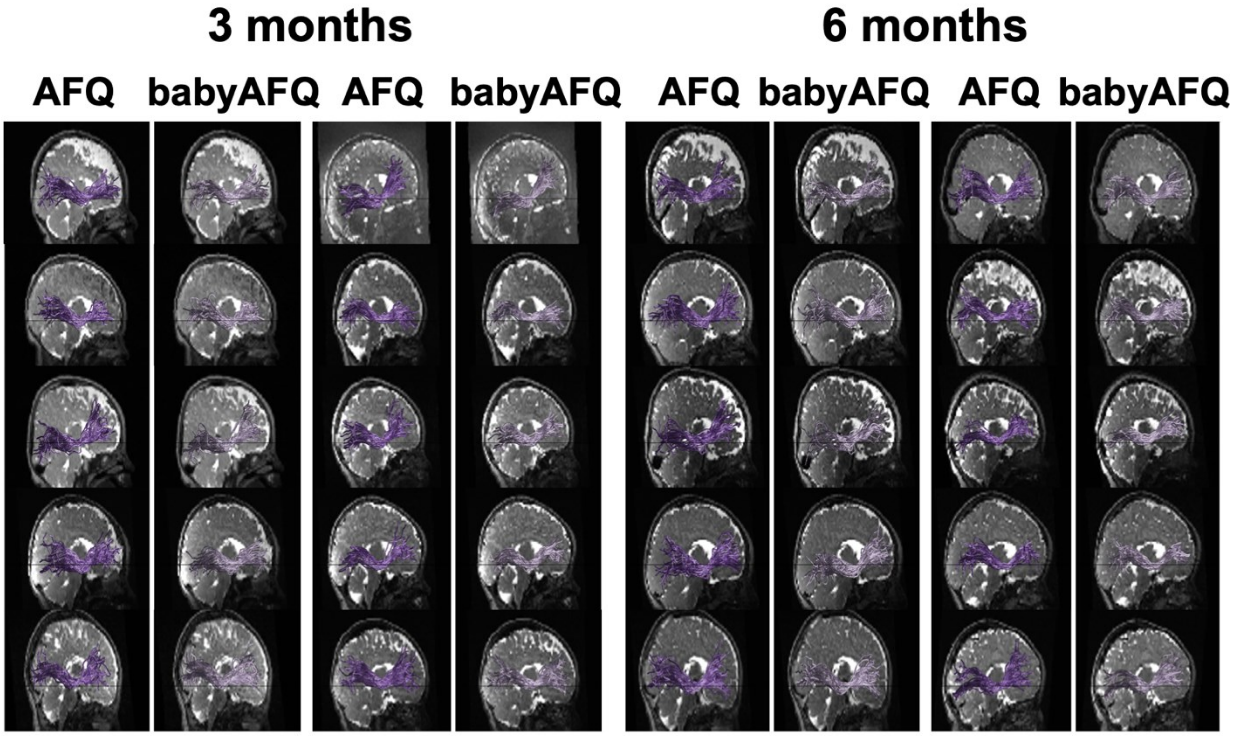

#### Left Inferior Longitudinal Fasciculus
#### Newborn

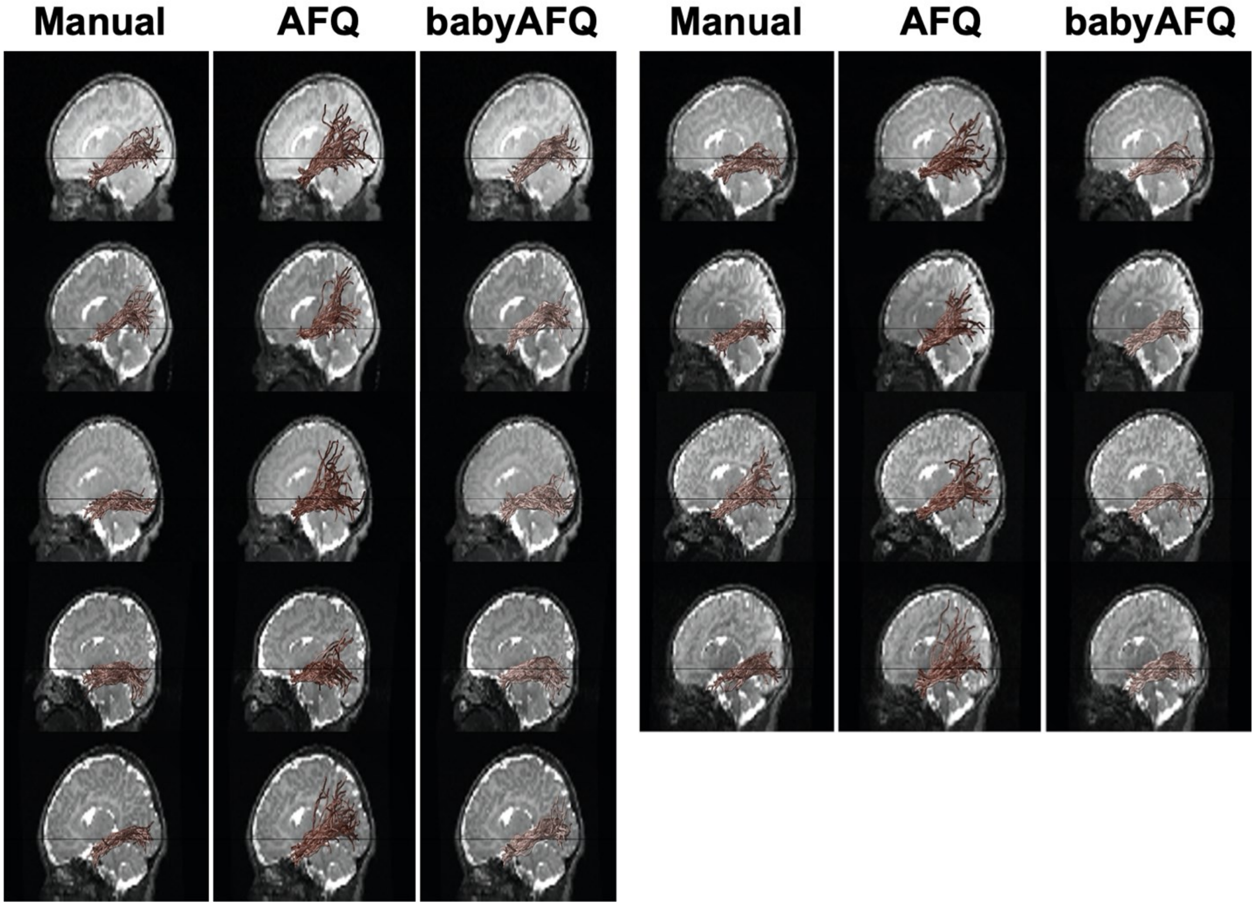

#### Left Inferior Longitudinal Fasciculus

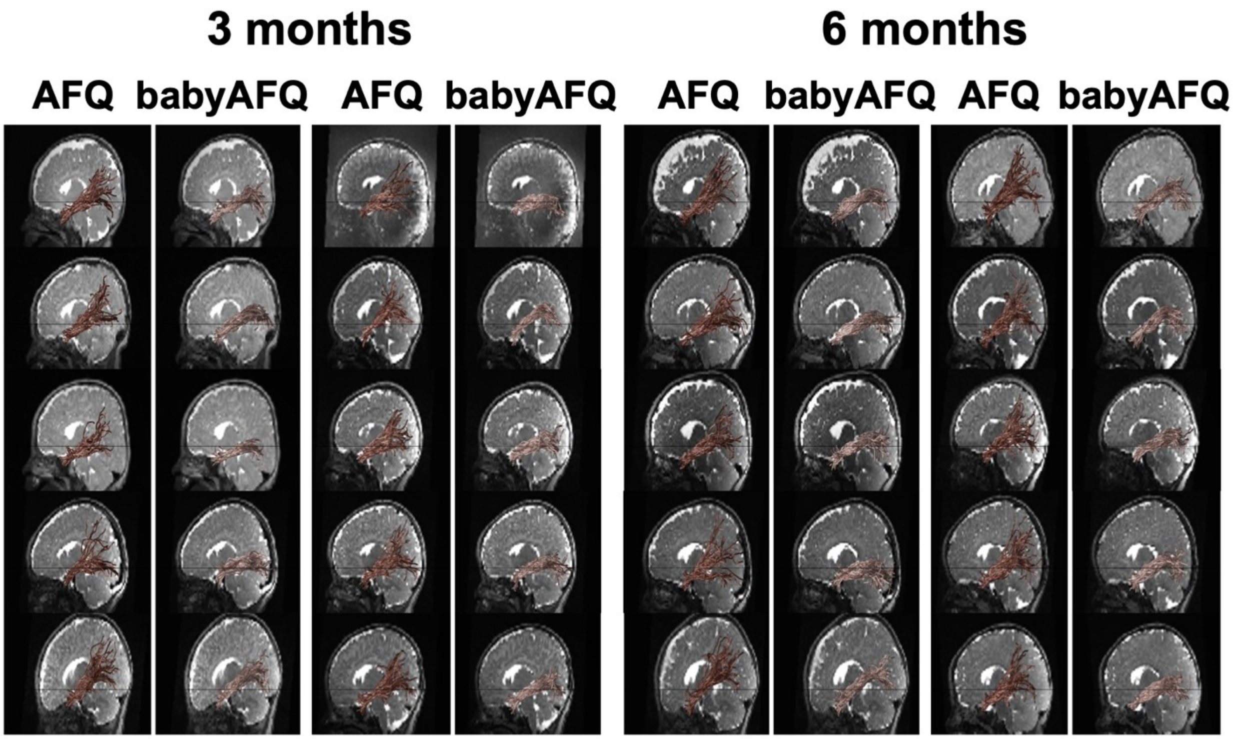

#### Right Inferior Longitudinal Fasciculus
#### Newborn

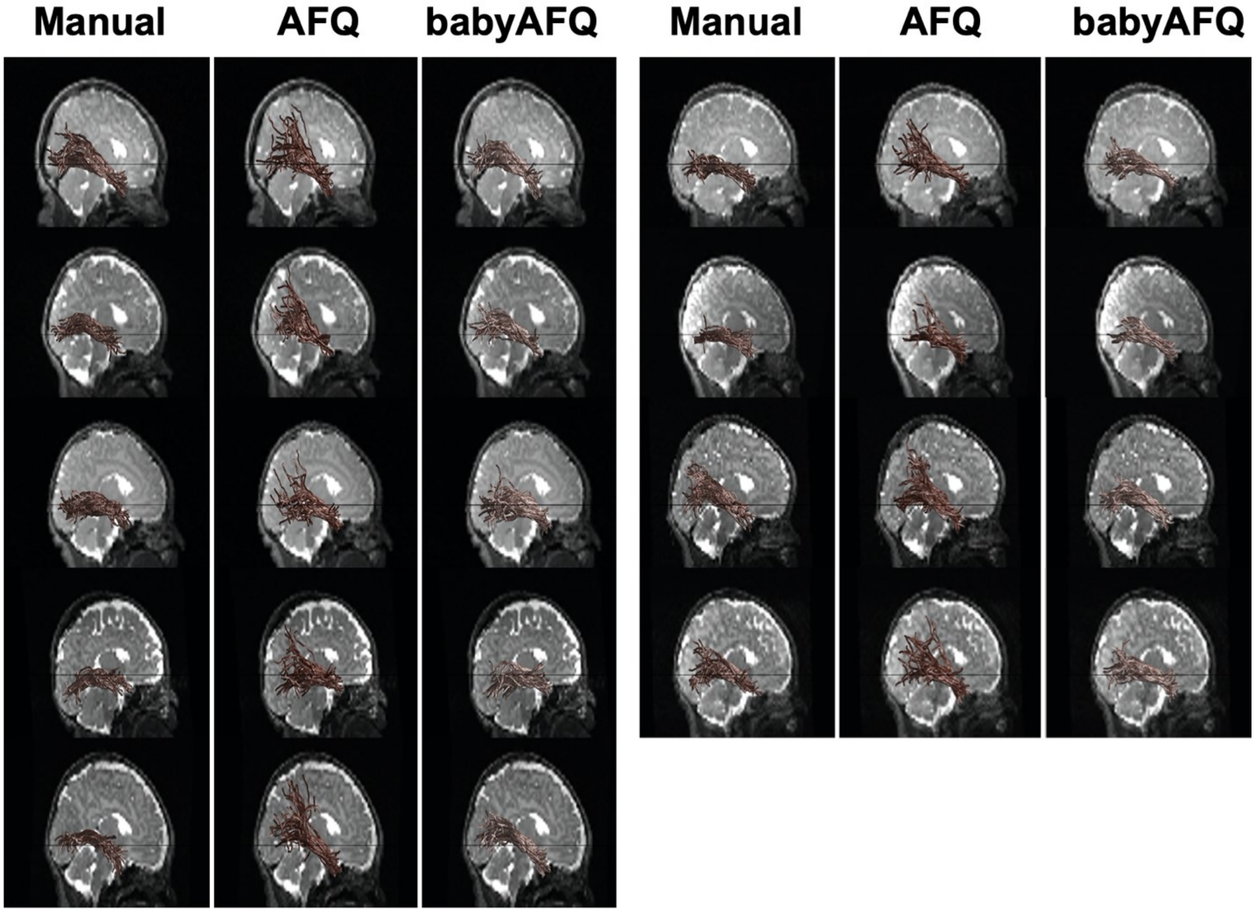

#### Right Inferior Longitudinal Fasciculus

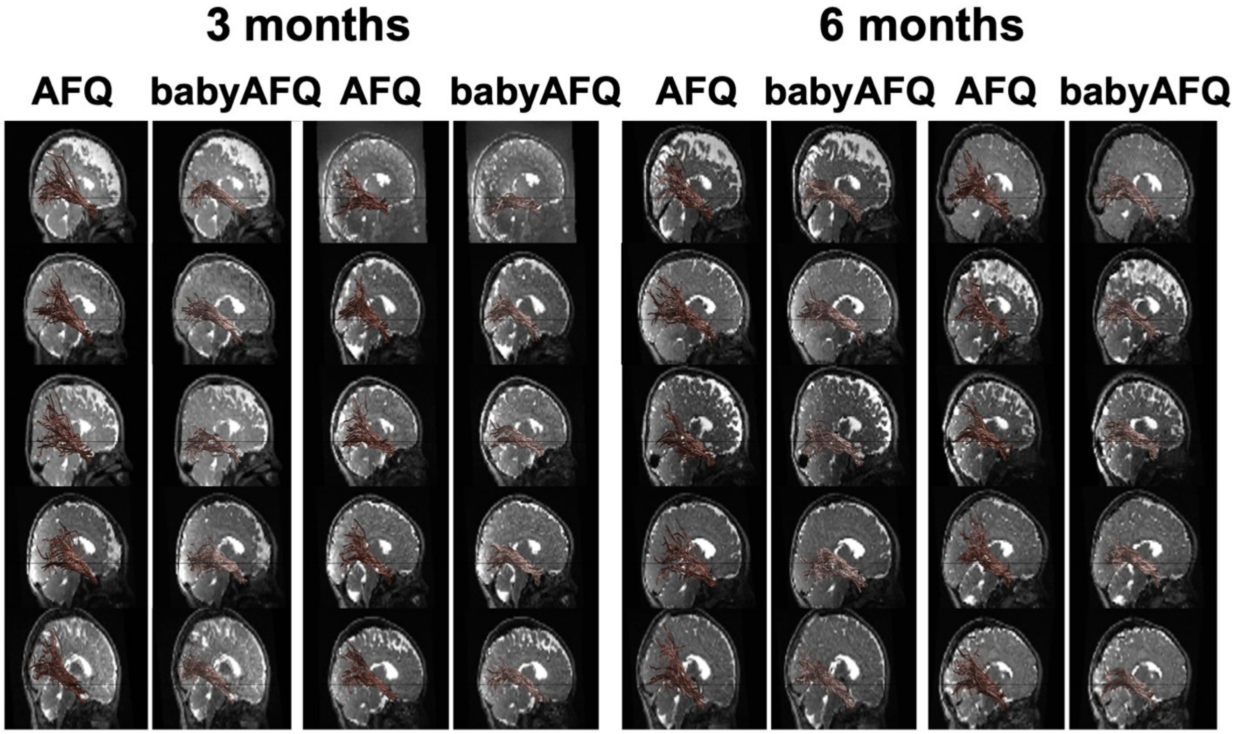

### Left Middle Longitudinal Fasciculus

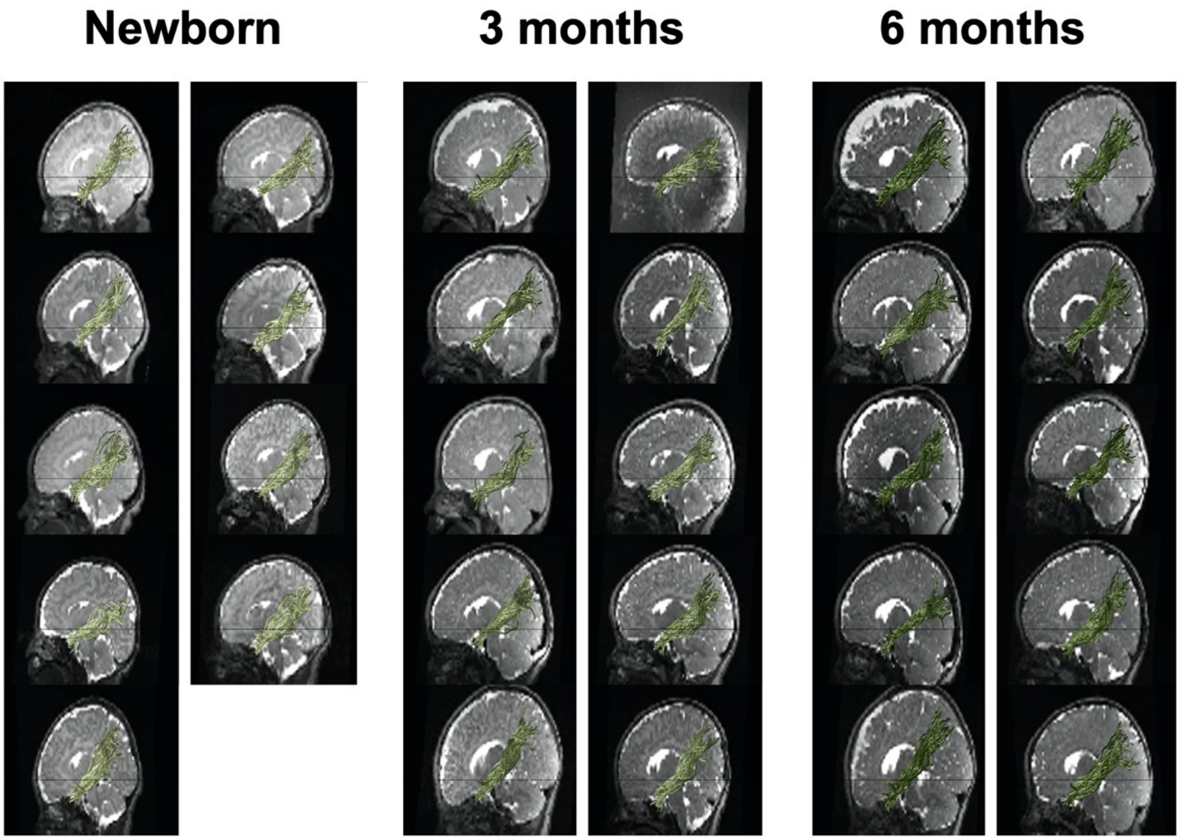

#### Right Middle Longitudinal Fasciculus

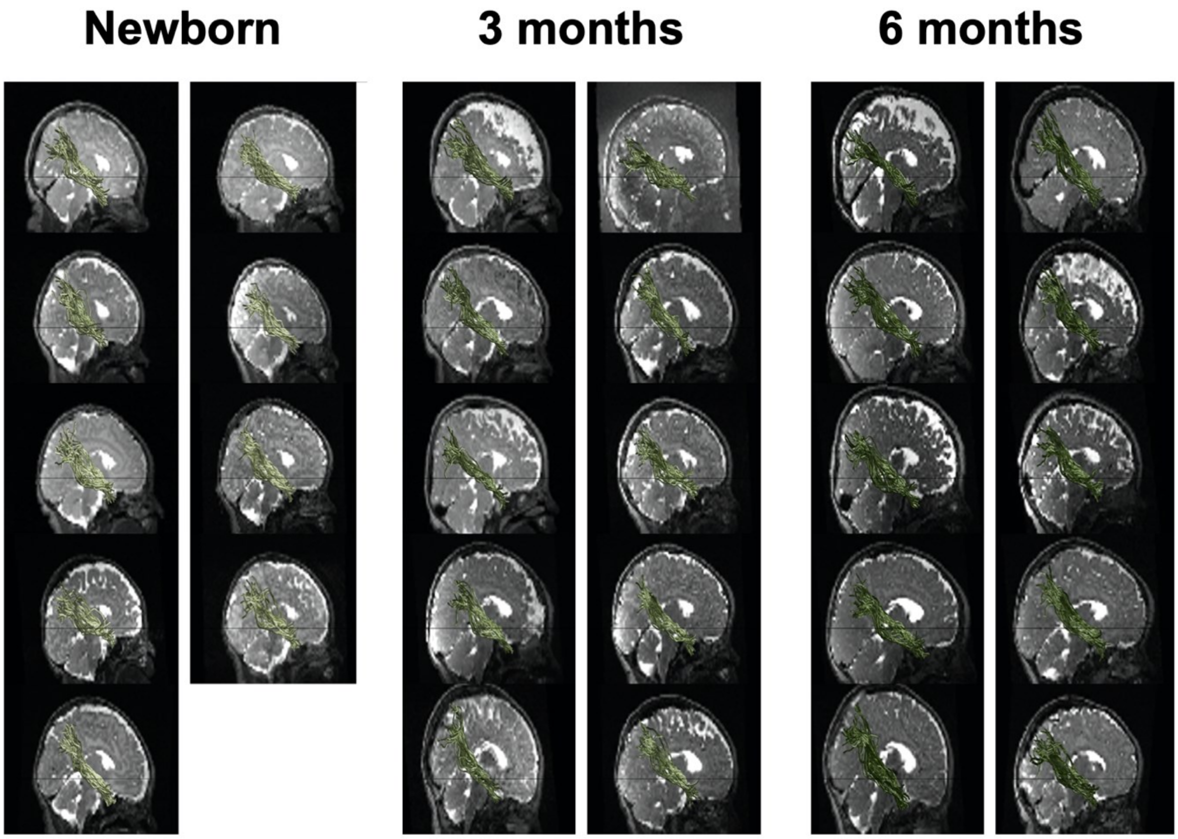

**Left Superior Longitudinal Fasciculus**
**Newborn**

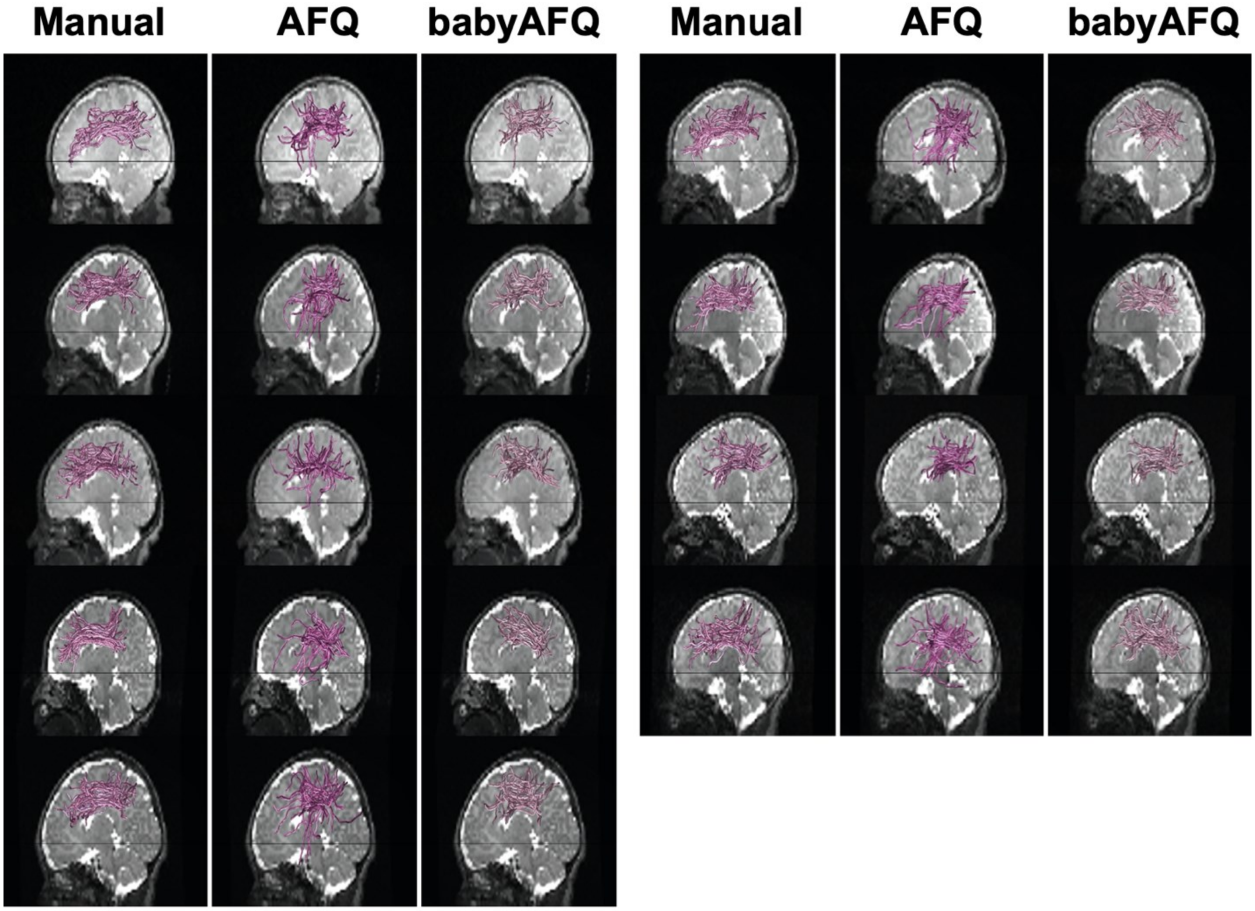

#### Left Superior Longitudinal Fasciculus

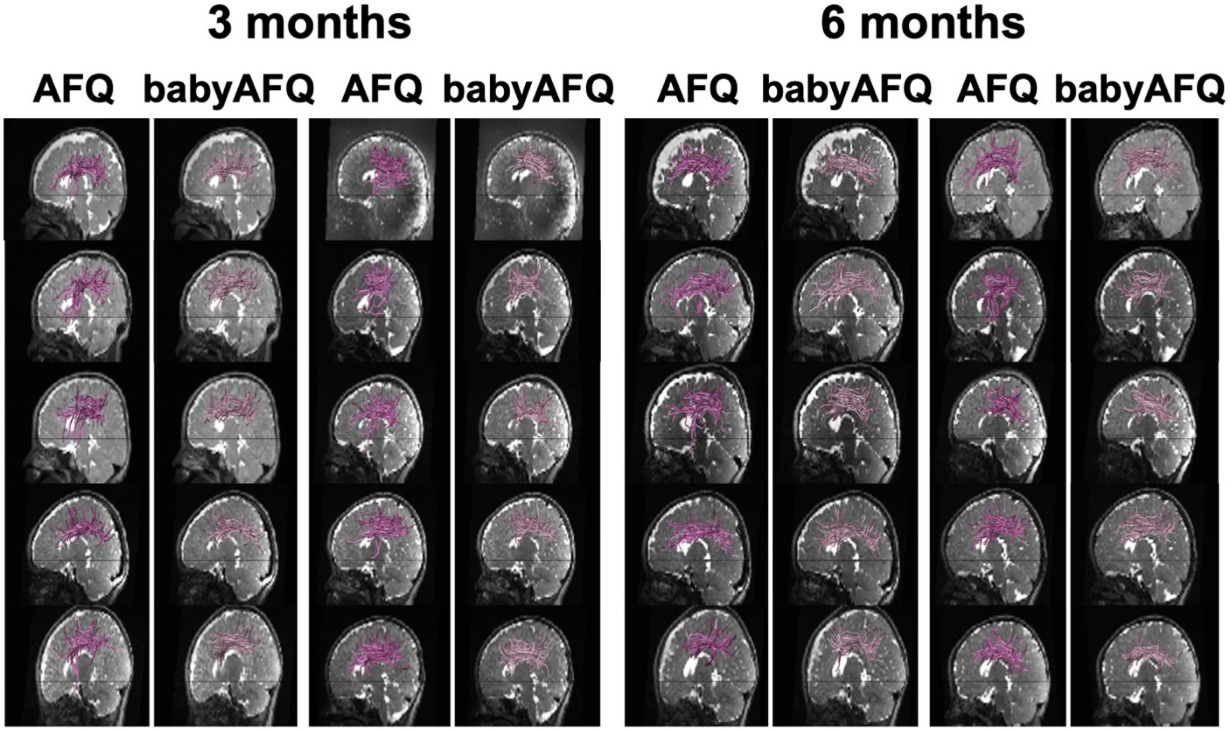

#### Right Superior Longitudinal Fasciculus
#### Newborn

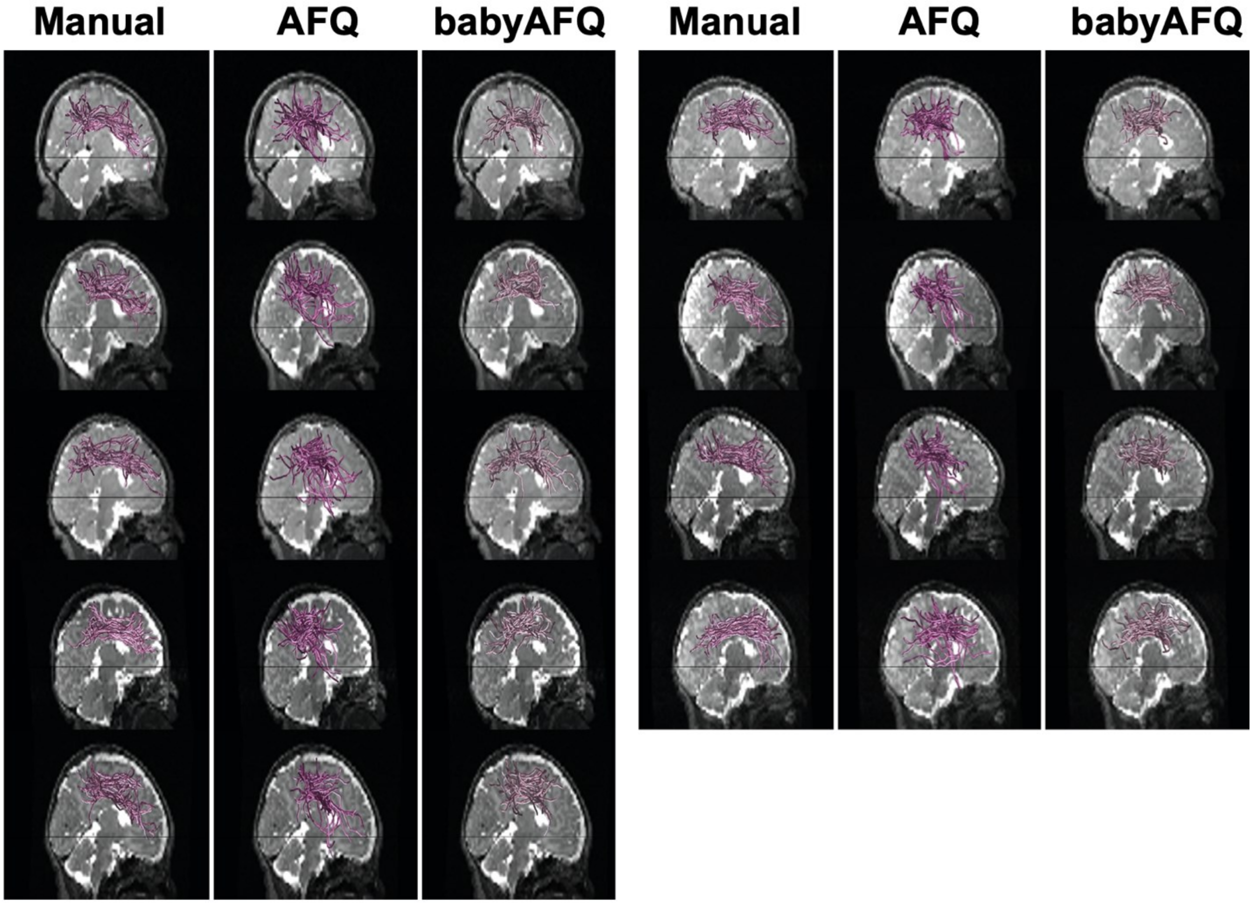

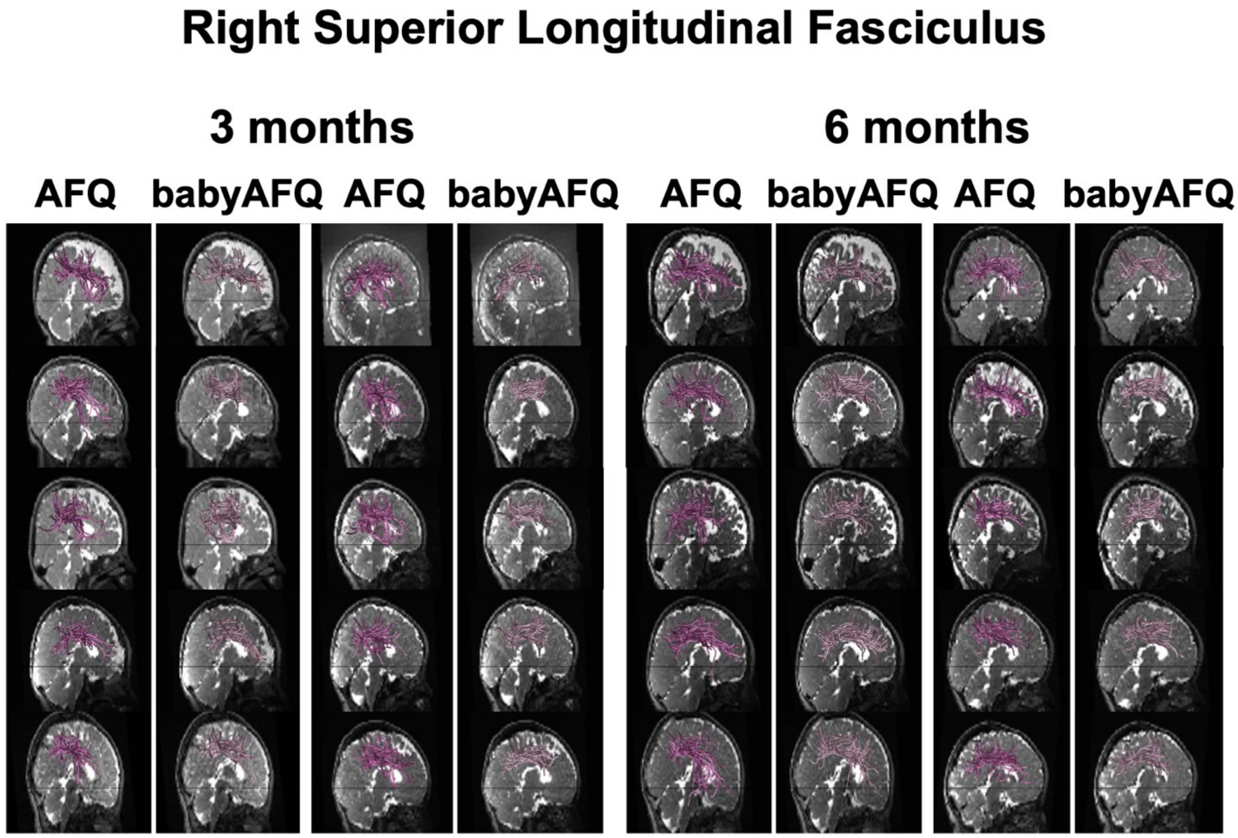

Right Superior Longitudinal Fasciculus
3 months
6 months
AFQ
babyAFQ
AFQ
babyAFQ
AFQ
babyAFQ
AFQ
babyAFQ

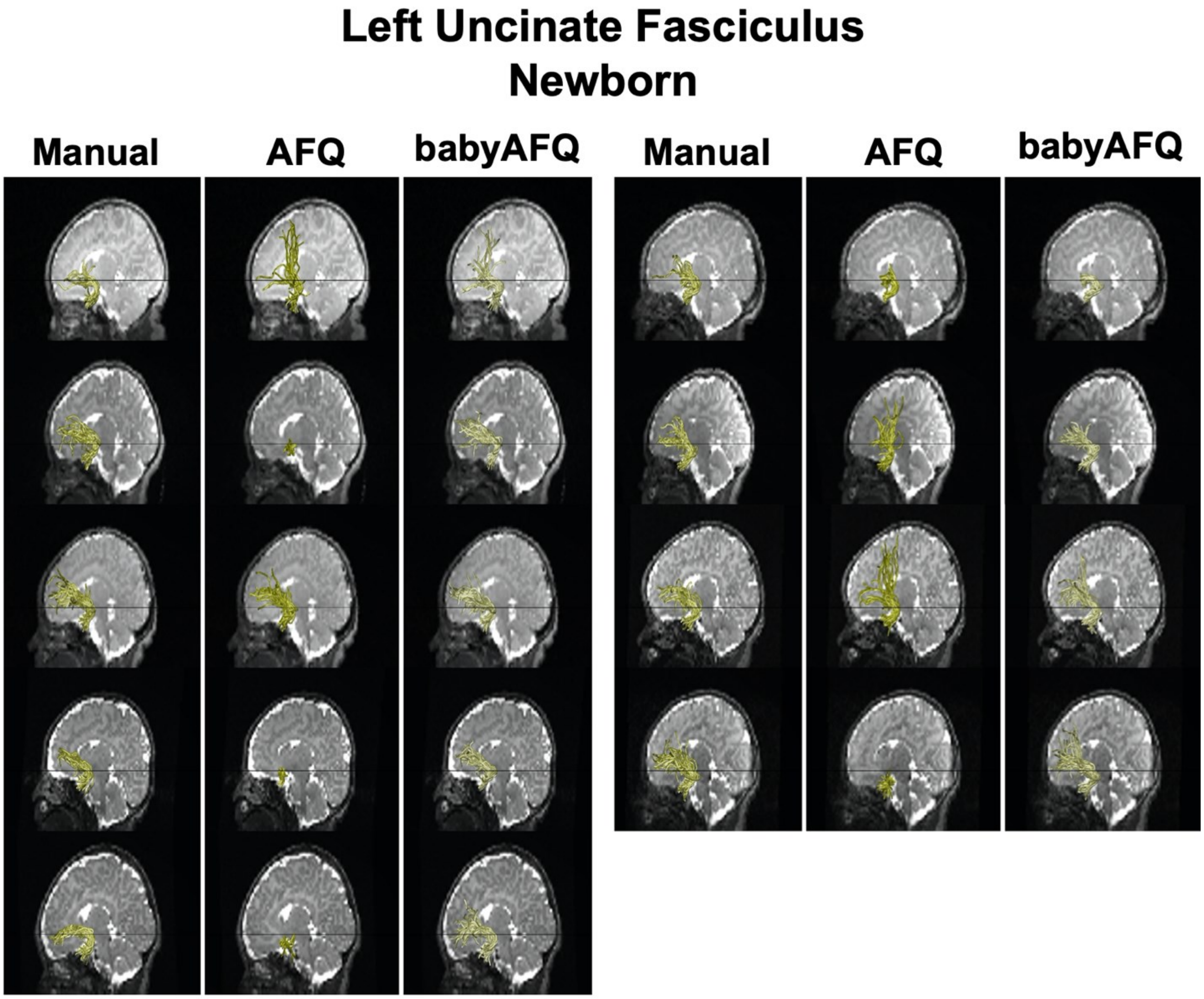
Left Uncinate Fasciculus
Newborn
Manual
AFQ
babyAFQ
Manual
AFQ
babyAFQ

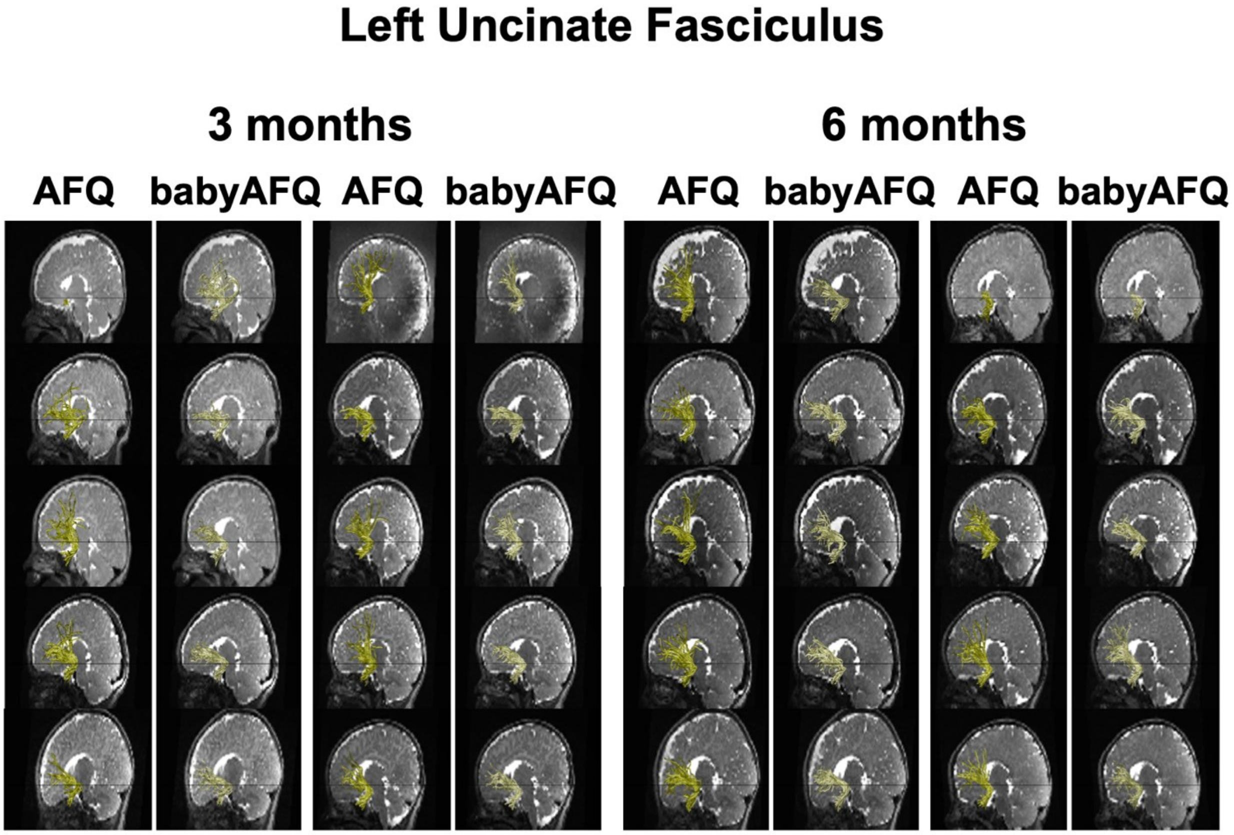
Left Uncinate Fasciculus
3 months
6 months
AFQ
babyAFQ
AFQ
babyAFQ
AFQ
babyAFQ
AFQ
babyAFQ

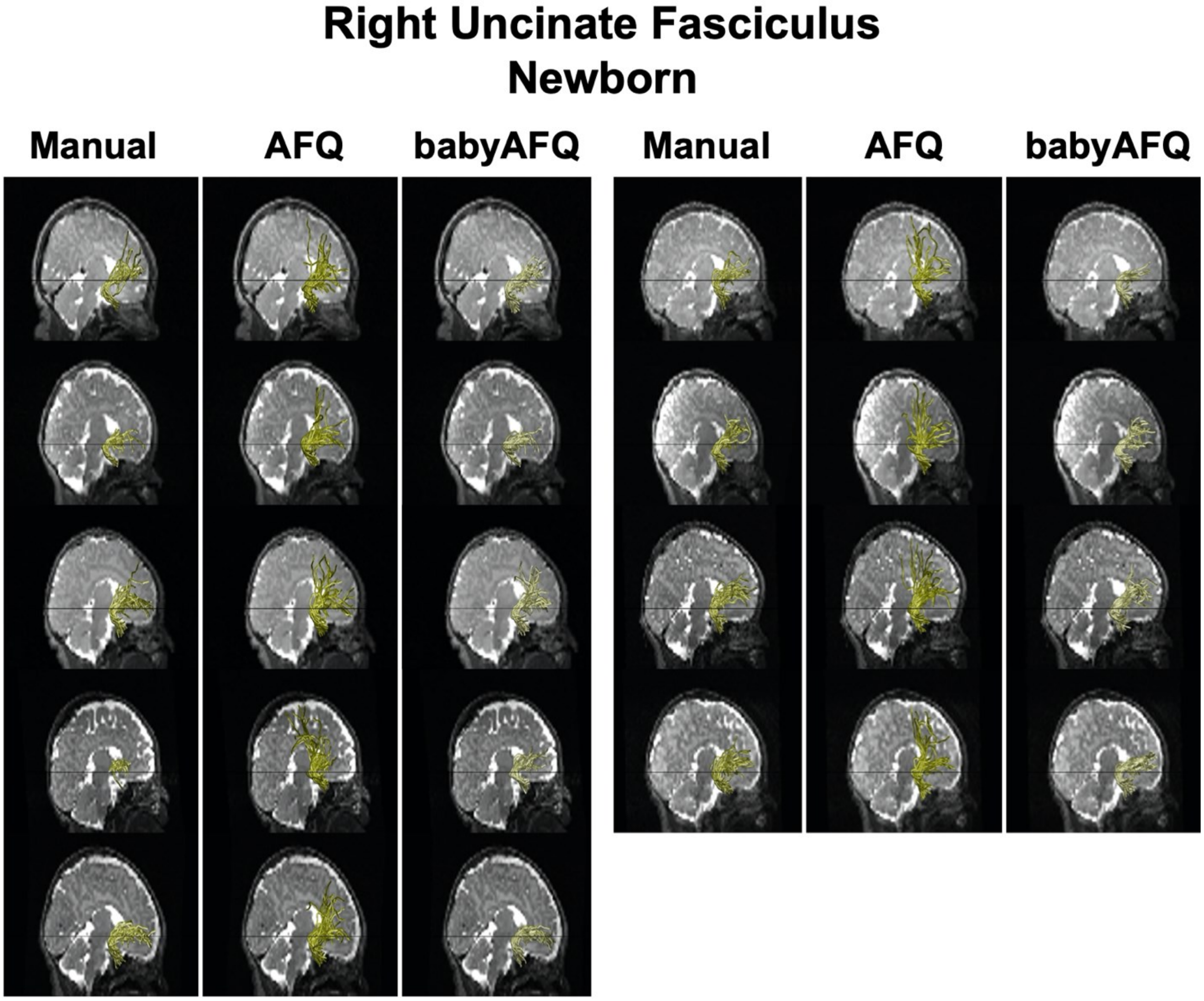
Right Uncinate Fasciculus
Newborn
Manual
AFQ
babyAFQ
Manual
AFQ
babyAFQ

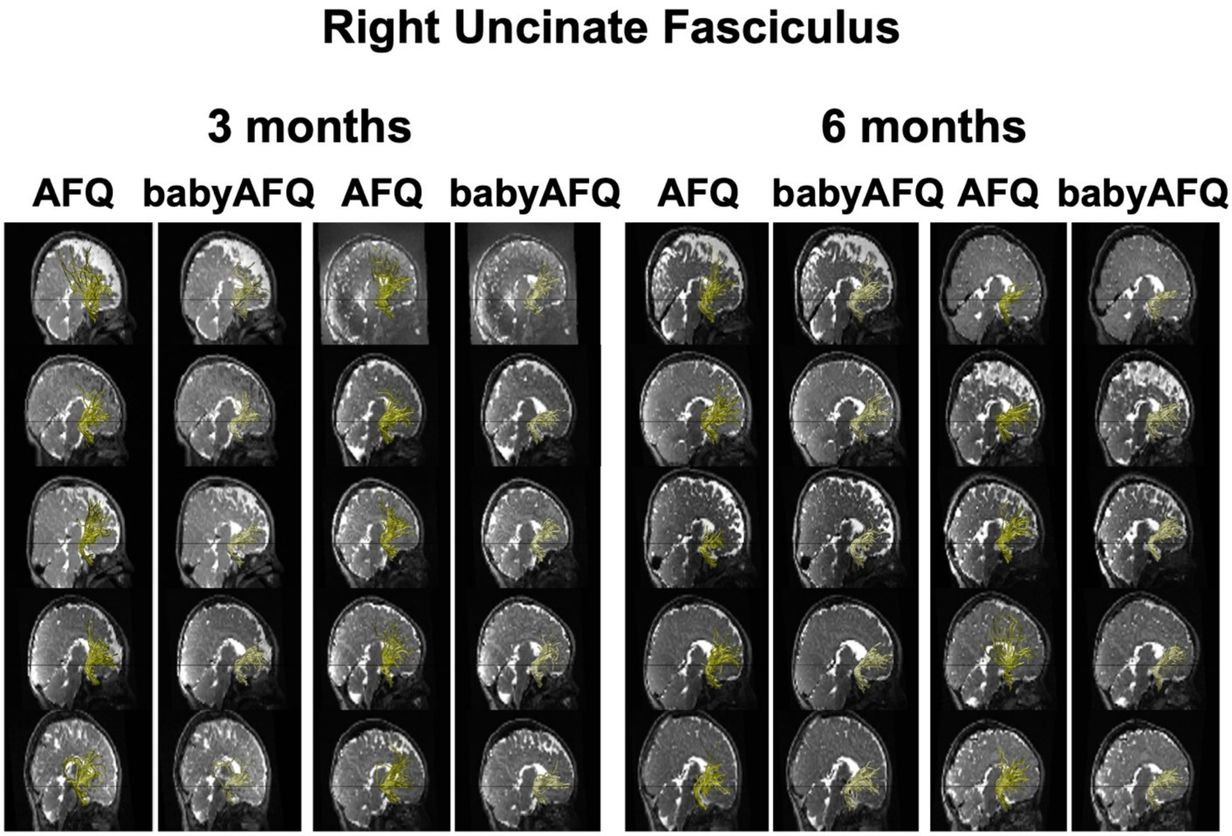
Right Uncinate Fasciculus
3 months
6 months
AFQ
babyAFQ
AFQ
babyAFQ
AFQ
babyAFQ
AFQ
babyAFQ

### Left Arcuate Fasciculus
#### Newborn

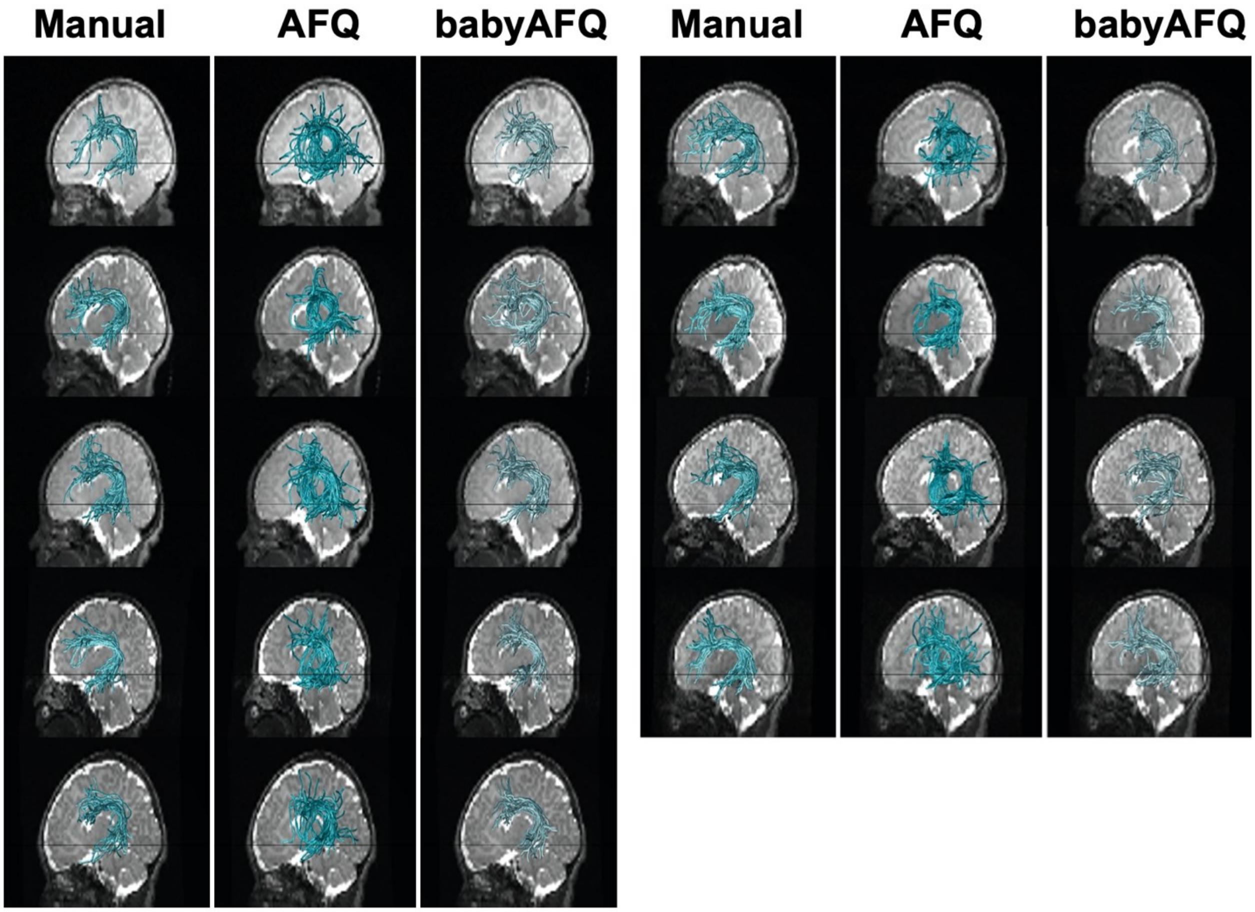

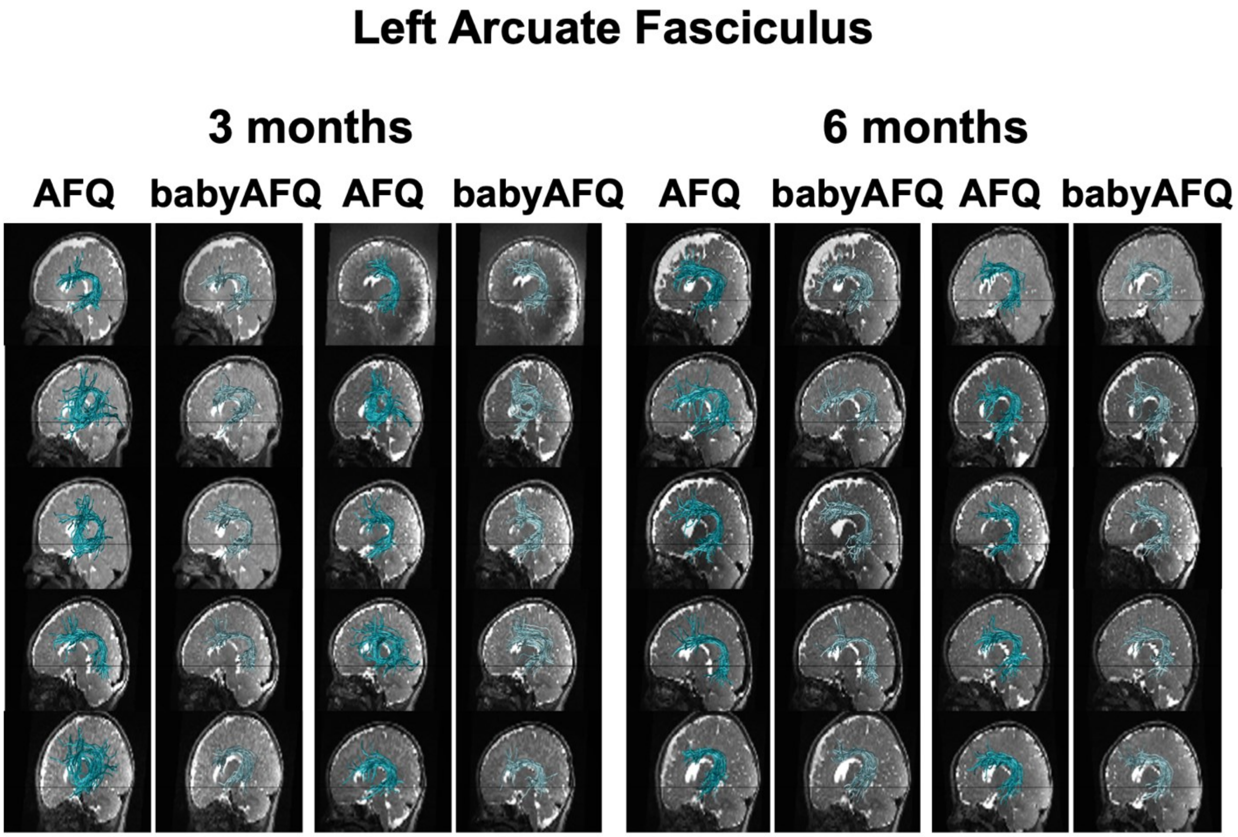
Left Arcuate Fasciculus
3 months
6 months
AFQ
babyAFQ
AFQ
babyAFQ
AFQ
babyAFQ
AFQ
babyAFQ

#### Right Arcuate Fasciculus
#### Newborn

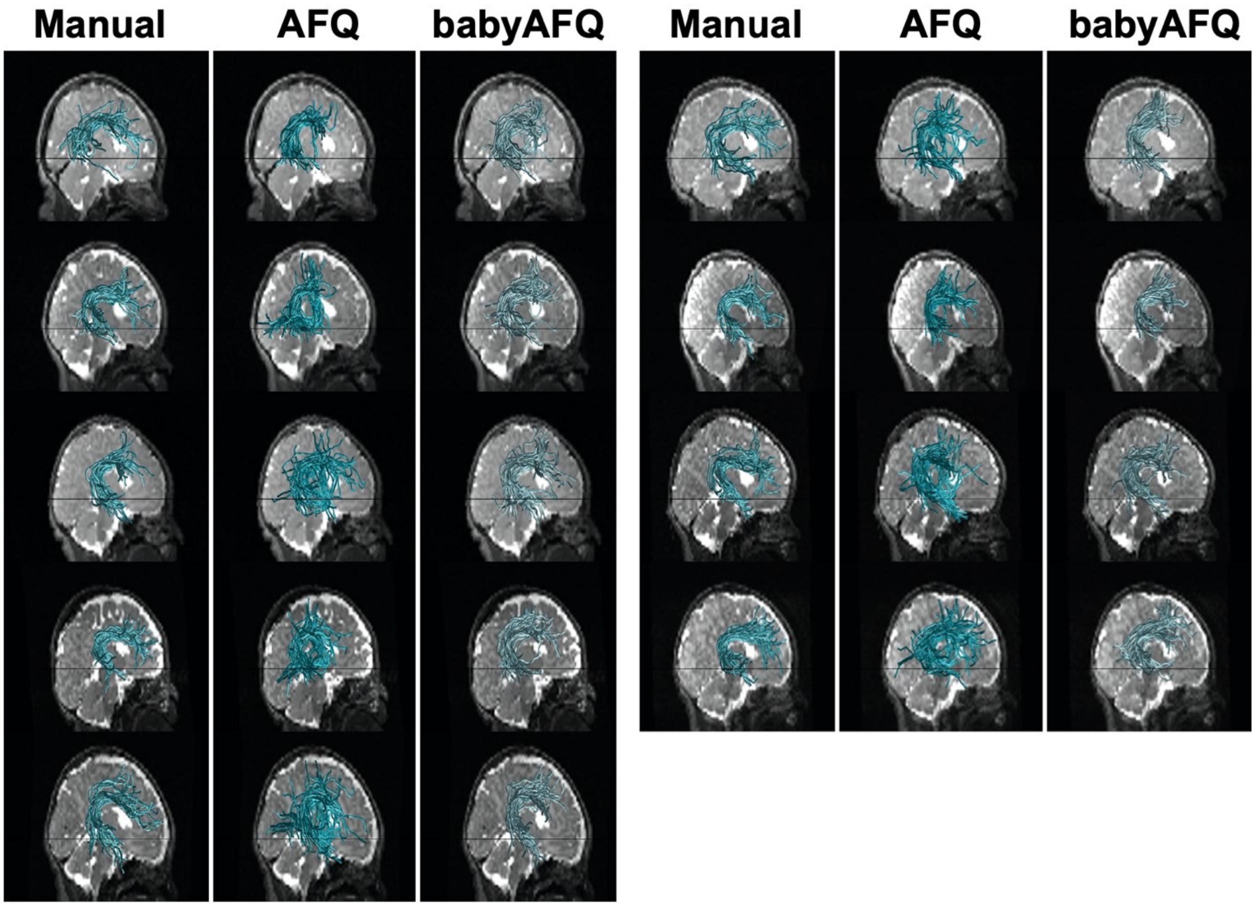

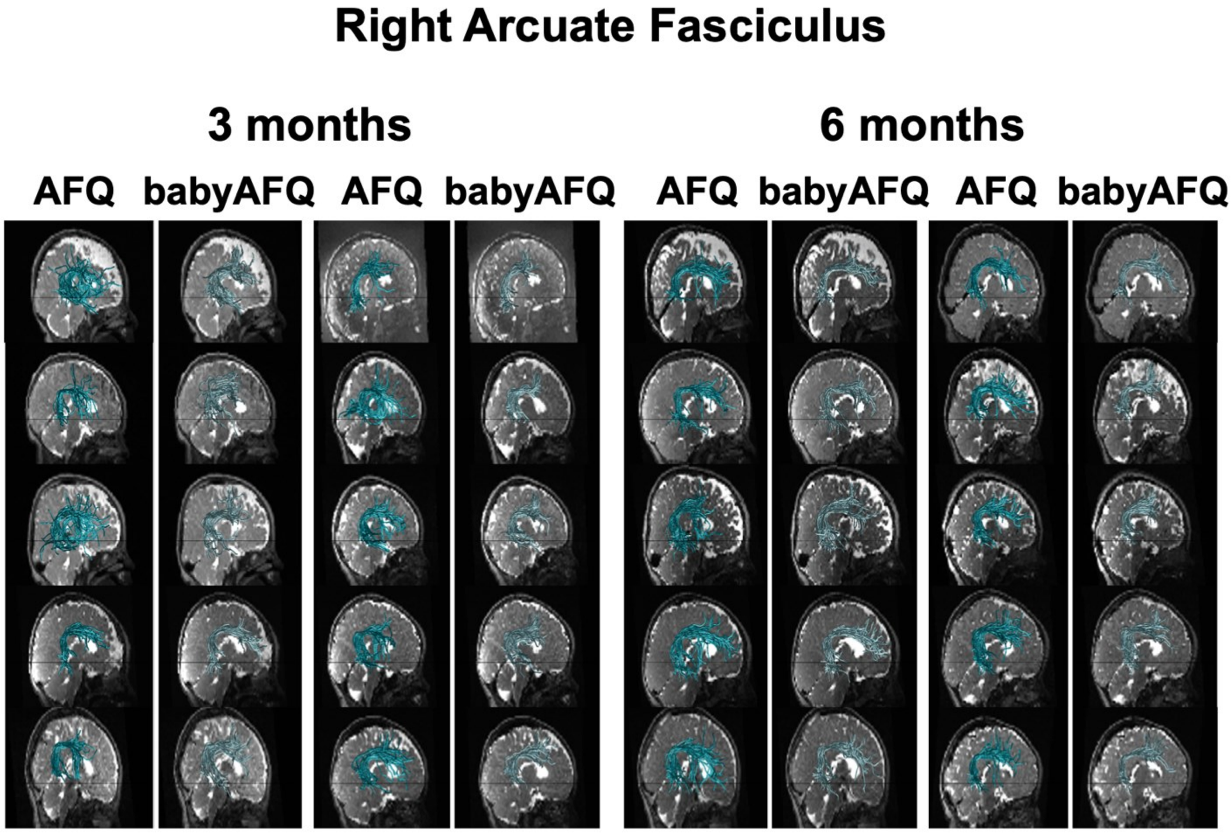
Right Arcuate Fasciculus
3 months
6 months
AFQ
babyAFQ
AFQ
babyAFQ
AFQ
babyAFQ
AFQ
babyAFQ

**Left Posterior Arcuate Fasciculus**
**Newborn**

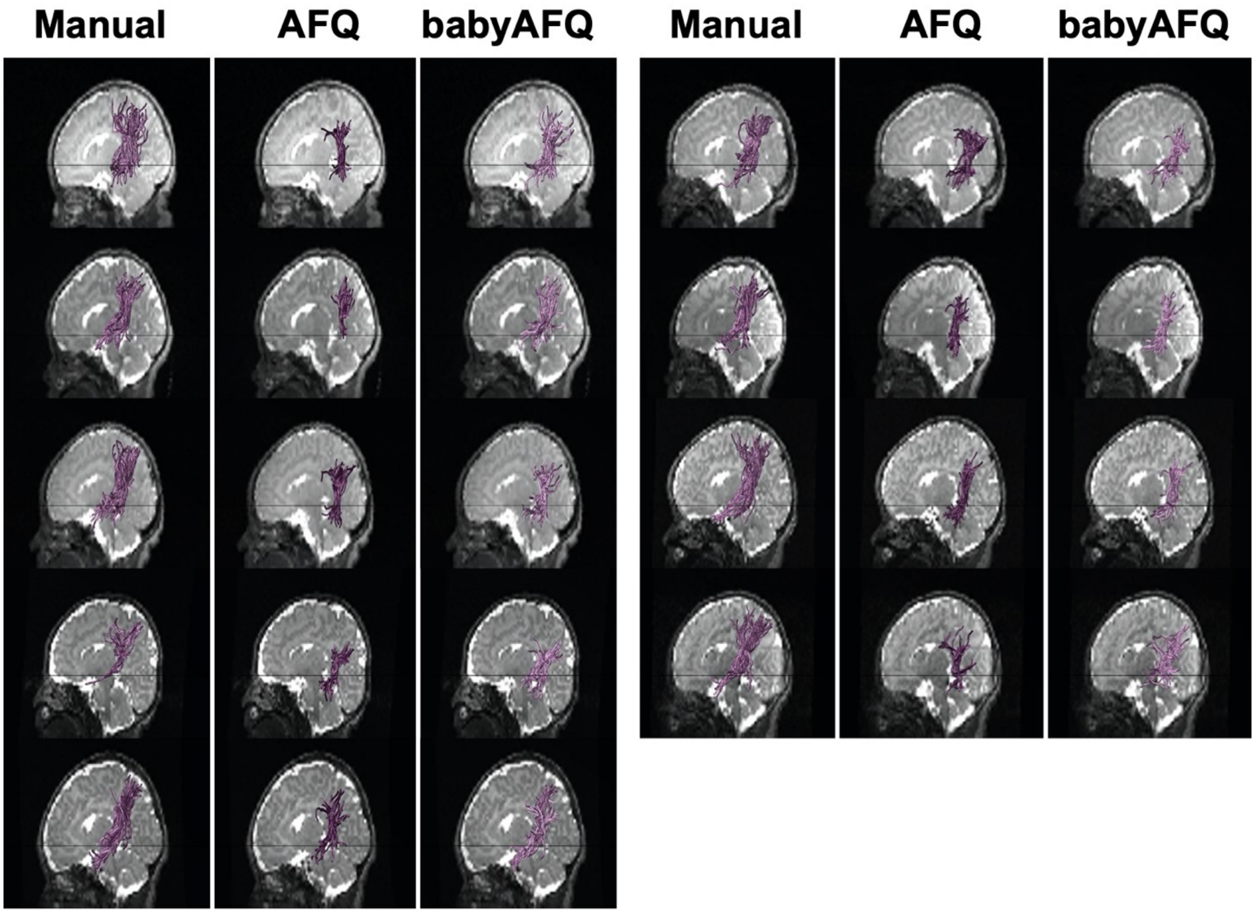

#### Left Posterior Arcuate Fasciculus

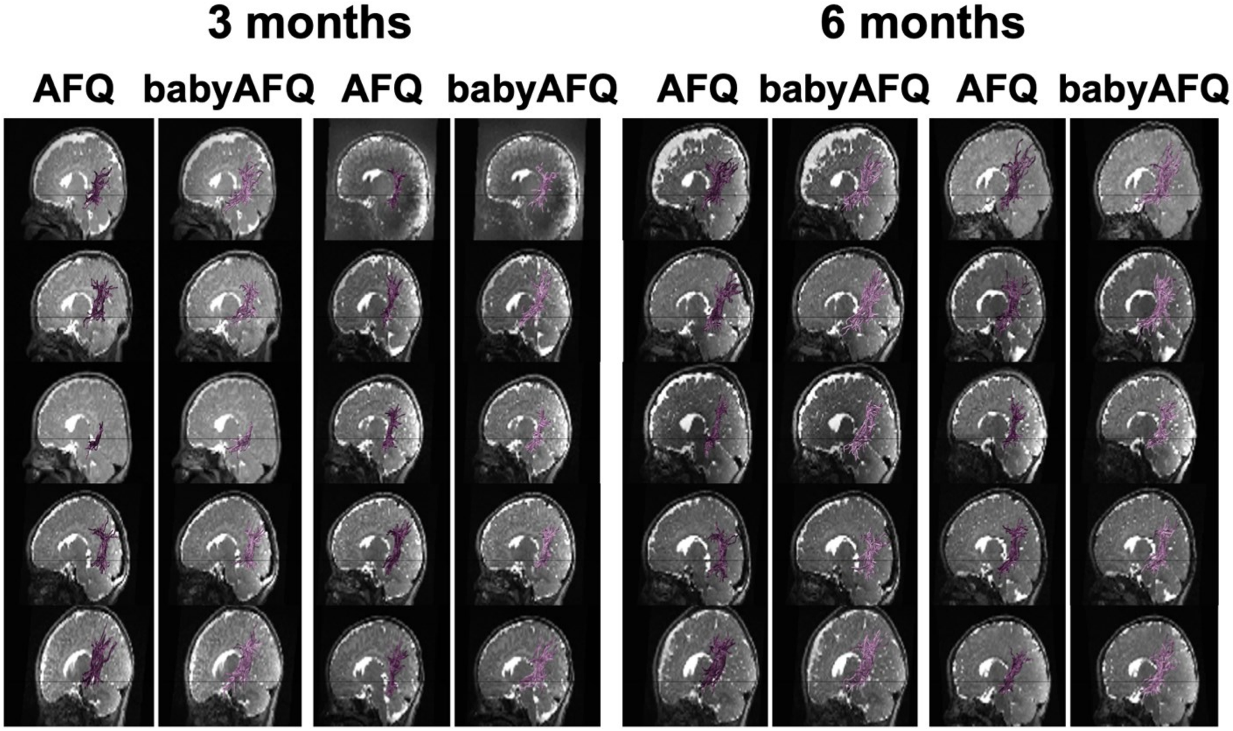

#### Right Posterior Arcuate Fasciculus
#### Newborn

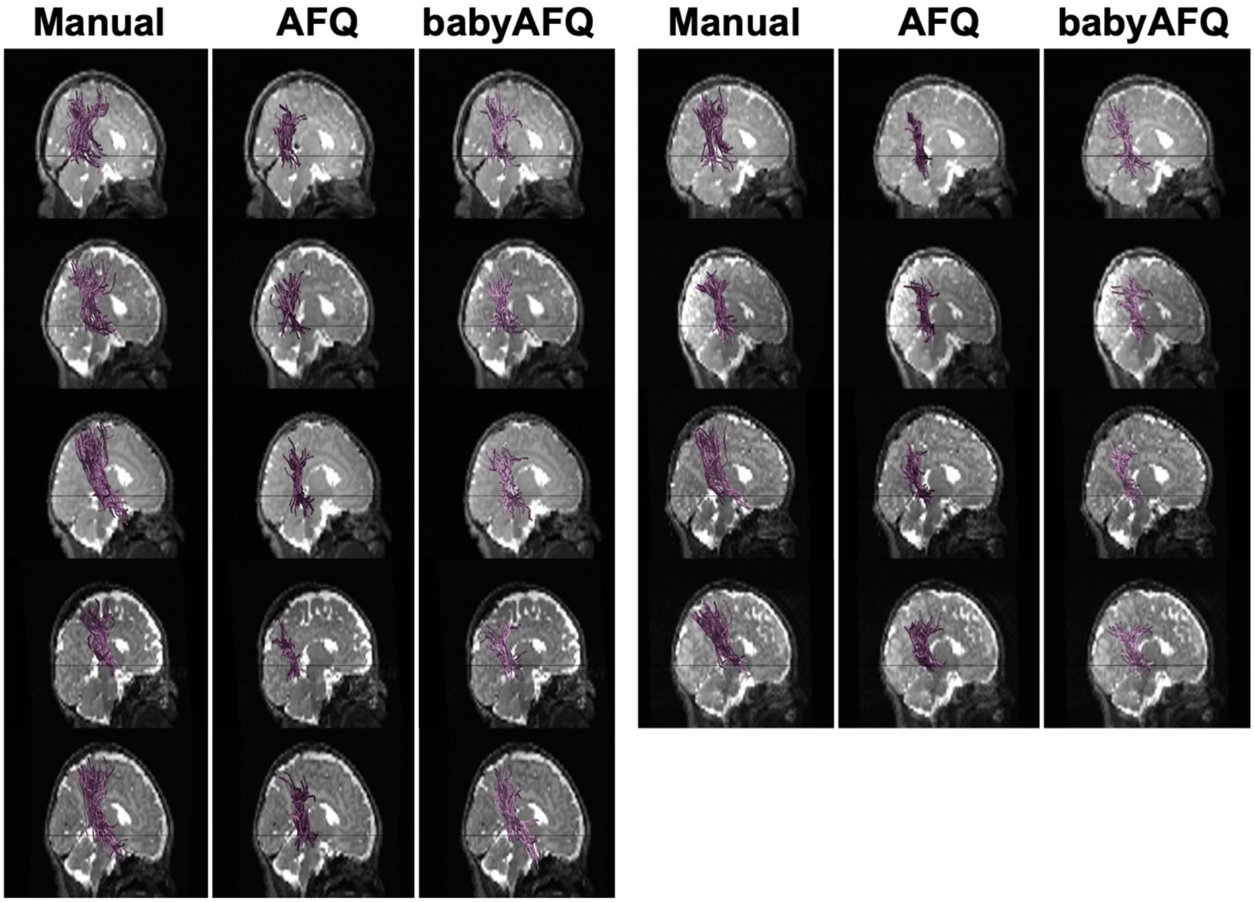

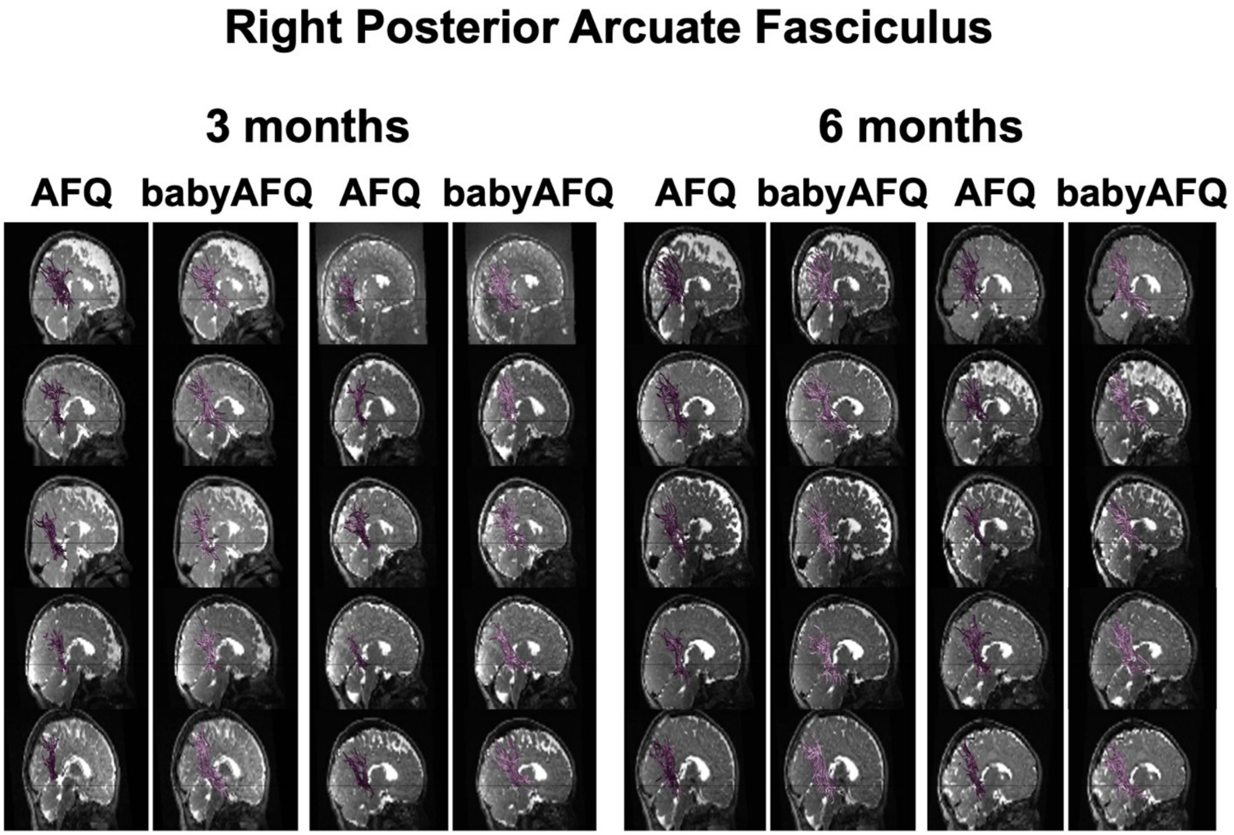
Right Posterior Arcuate Fasciculus
3 months
6 months
AFQ
babyAFQ
AFQ
babyAFQ
AFQ
babyAFQ
AFQ
babyAFQ

### Left Vertical Occipital Fasciculus
#### Newborn

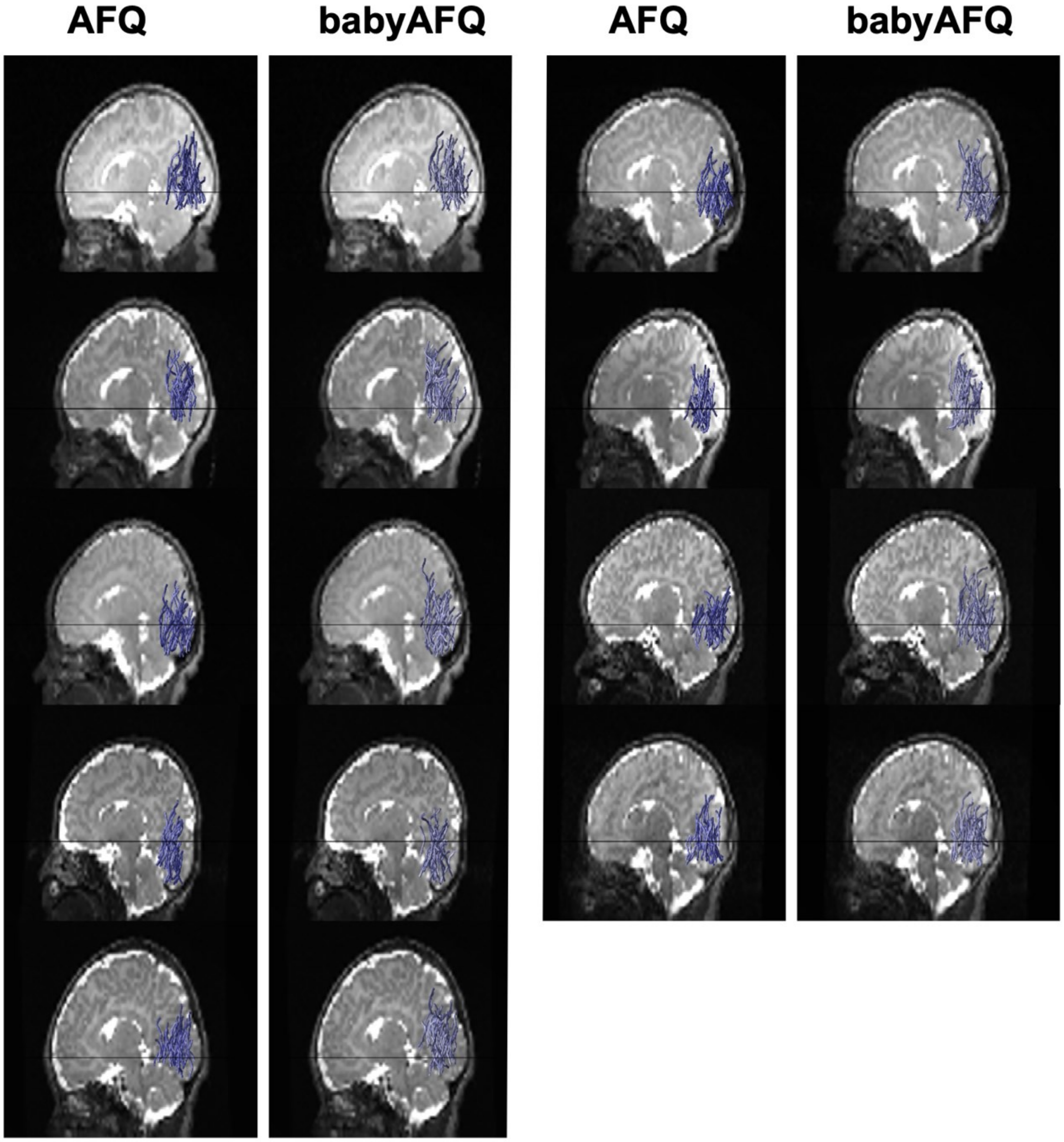

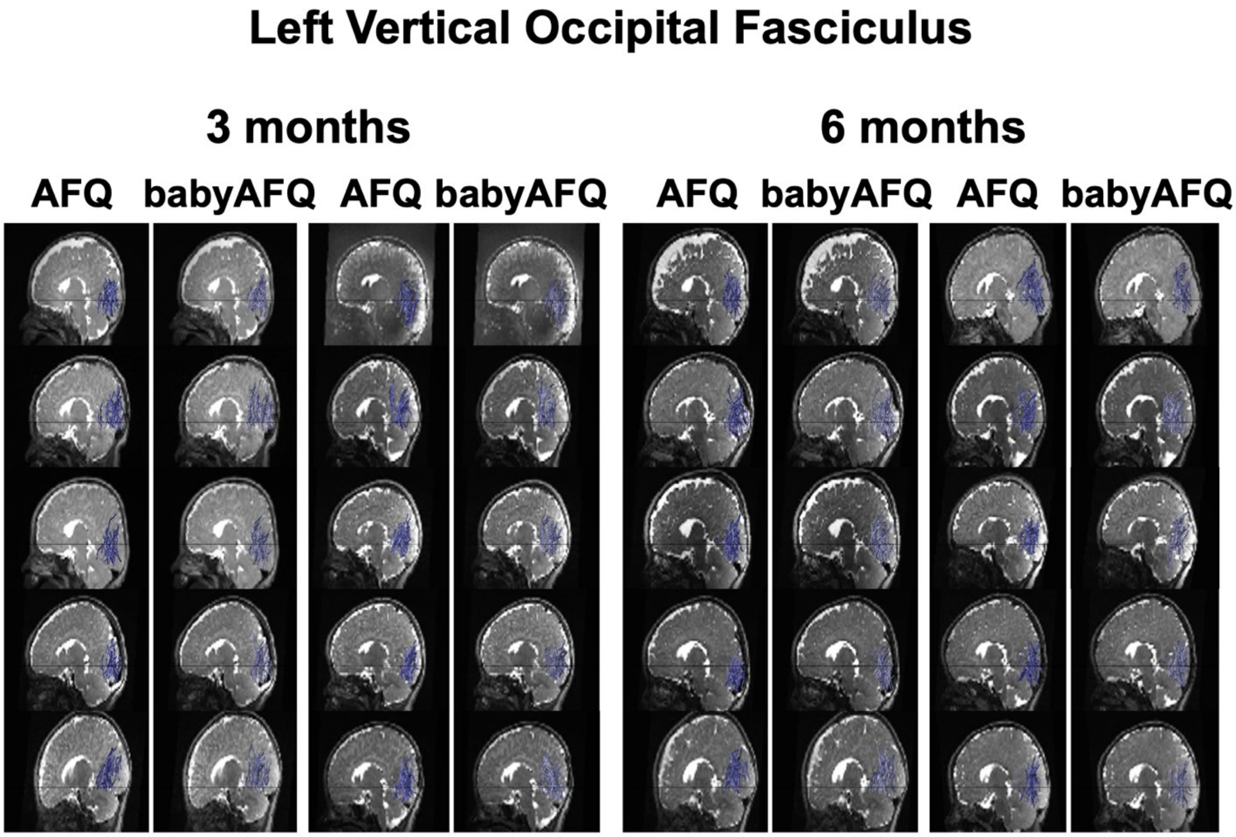
Left Vertical Occipital Fasciculus
3 months
6 months
AFQ
babyAFQ
AFQ
babyAFQ
AFQ
babyAFQ
AFQ
babyAFQ

### Right Vertical Occipital Fasciculus
#### Newborn

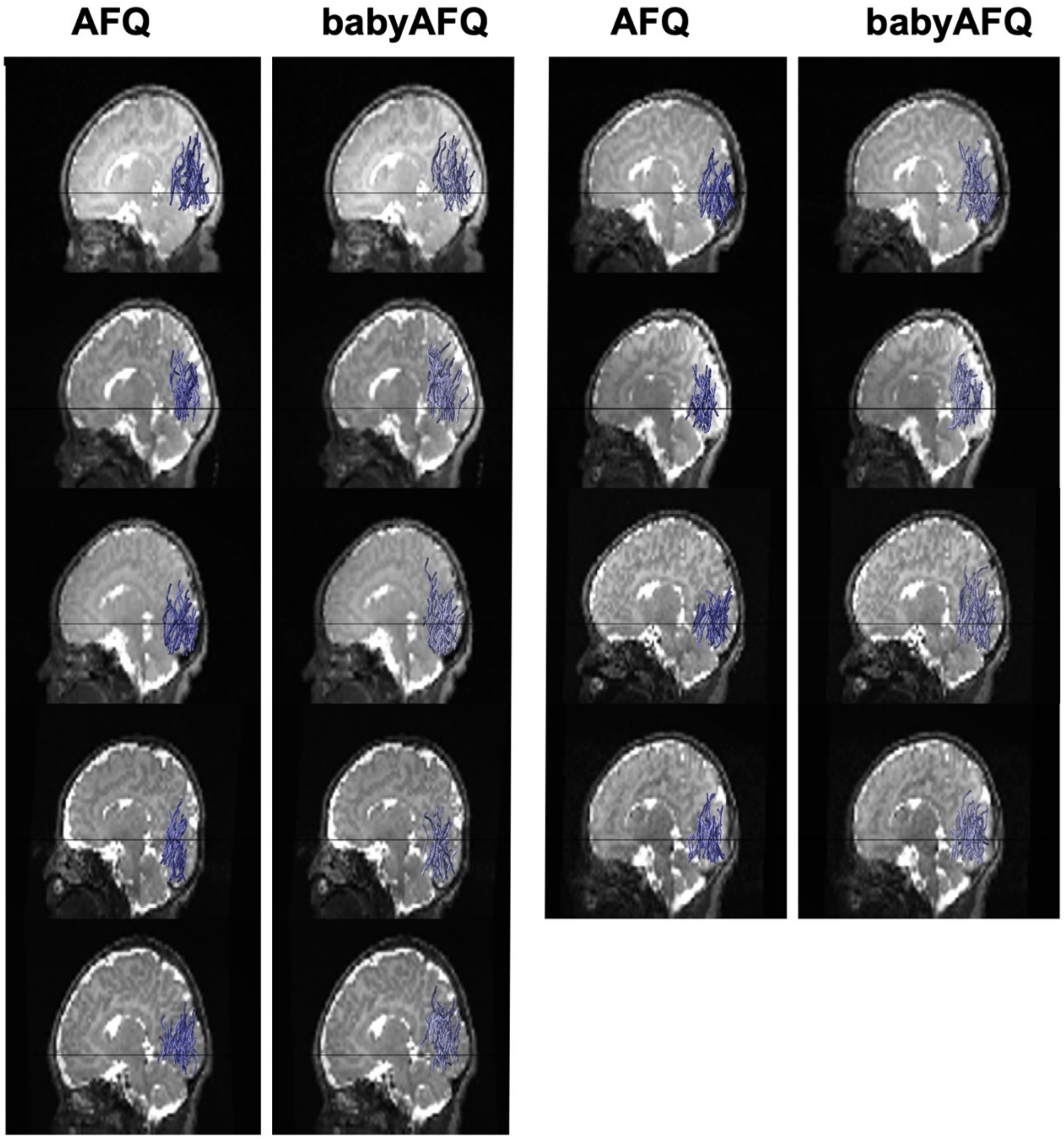

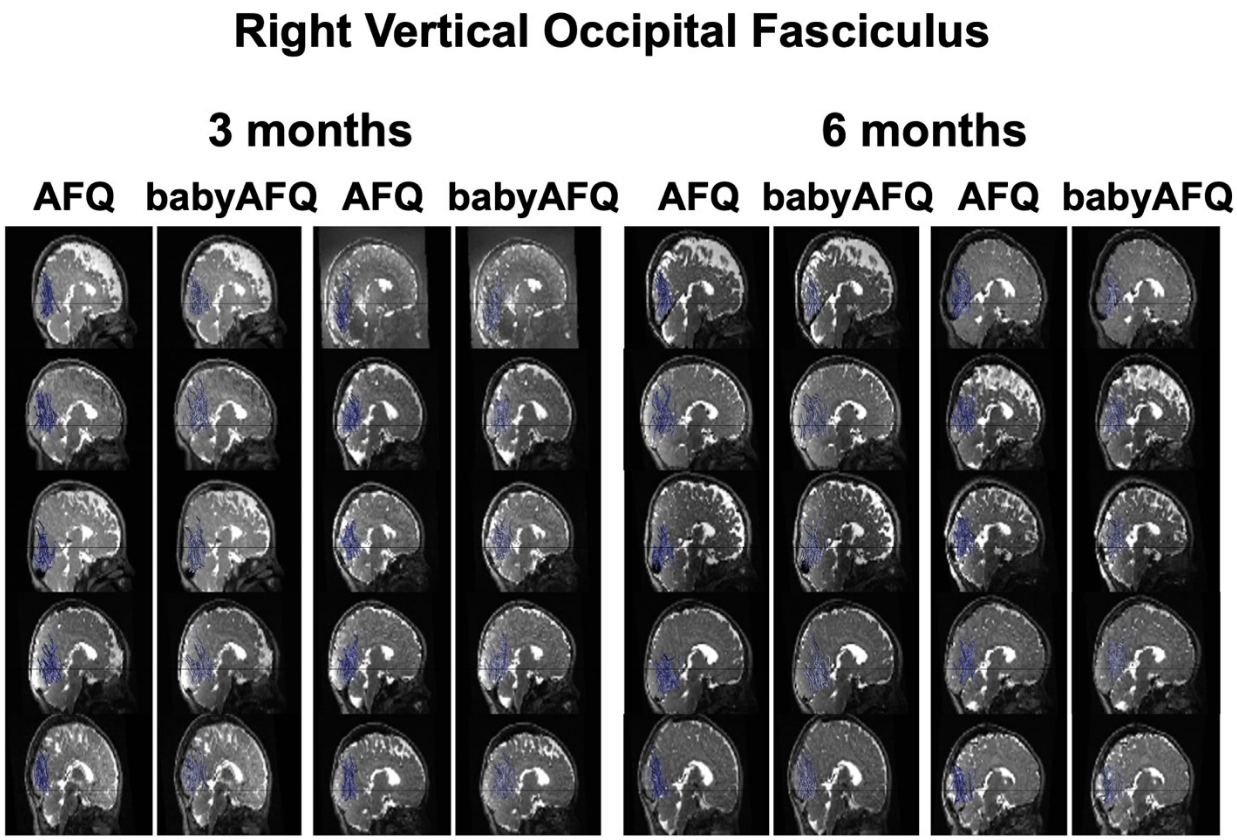
Right Vertical Occipital Fasciculus
3 months
6 months
AFQ
babyAFQ
AFQ
babyAFQ
AFQ
babyAFQ
AFQ
babyAFQ
